# Extreme intratumour heterogeneity and driver evolution in mismatch repair deficient gastro-oesophageal cancer

**DOI:** 10.1101/755199

**Authors:** Katharina von Loga, Andrew Woolston, Marco Punta, Louise Barber, Beatrice Griffiths, Maria Semiannikova, Georgia Spain, Benjamin Challoner, Kerry Fenwick, Ronald Simon, Andreas Marx, Guido Sauter, Stefano Lise, Nik Matthews, Marco Gerlinger

## Abstract

Mismatch repair deficient (dMMR) gastro-oesophageal adenocarcinomas (GOAs) show better outcomes than their MMR-proficient counterparts and high immunotherapy sensitivity. The hypermutator-phenotype of dMMR tumours theoretically enables high evolvability but how these cancers evolve has not been investigated in detail. We applied multi-region exome sequencing (MSeq) to four treatment-naïve dMMR GOAs. This revealed extreme intratumour heterogeneity (ITH), exceeding ITH in other cancer types >20-fold, but also long phylogenetic trunks which may explain the exquisite immunotherapy sensitivity of dMMR tumours. Subclonal driver mutations were common and parallel evolution occurred in *RAS, PIK3CA*, SWI/SNF-complex genes and in immune evasion regulators. MSeq data and evolution analysis of single region-data from 64 MSI GOAs showed that chromosome 8 gains were among the earliest genetic events and that the hypermutator-phenotype remained active during progression. MSeq may be necessary for biomarker development in these heterogeneous cancers. Comparison with other MSeq-analysed tumour types revealed mutation rates and their timing as major determinants of phylogenetic tree morphologies.

## Introduction

Gastro-oesophageal adenocarcinomas (GOAs) are one of the commonest causes of cancer related mortality worldwide^1^. Microsatellite instable (MSI) and DNA mismatch repair deficient (dMMR) cancers are a distinct subtype of GOAs with a reported prevalence of up to ∼20%^2-4^ in the stomach and gastro-oesophageal junction but which are rare in the oesophagus^5,6^. dMMR results from genetic inactivation of *MLH1, MSH2, MSH6* or *PMS2* or methylation of *MLH1*. These tumours are characterized by a hypermutator phenotype leading to high mutation loads and a large fraction of small insertions and deletions (indels), predominantly in homopolymer repeats and dinucleotide repeats termed microsatellites. dMMR GOAs have distinct clinical characteristics compared to their MMR proficient counterparts, including lower TNM-stage at presentation and better survival^3^. This has been attributed to a large number of neoantigens encoded by the high mutation burden which promotes immune-surveillance by the adaptive immune system. Consistent with the notion of high immunogenicity, dMMR cancers are among the solid tumour types most sensitive to checkpoint-inhibiting immunotherapy with response rates of up to 85.7% reported in small series^7,8^. However, not all of these tumours respond to immunotherapy and some acquire resistance after initial responses. Chemotherapy and anti-angiogenic drugs are the only other systemic treatment options for dMMR GOAs and the identification of novel therapeutics is important to improve outcomes.

Genetic intratumour heterogeneity (ITH) and ongoing cancer evolution that leads to the acquisition of novel driver mutations have been demonstrated in multiple cancer types^9^. The ability to evolve is thought to foster cancer progression, drug resistance and poor outcomes in cancer medicine. High mutation rates may fuel cancer evolvability by generating an abundance of novel phenotypes which selection can act upon^10^. A pan-cancer study indeed demonstrated large numbers of subclonal mutations within single tumour regions of MSI cancers^11^. However, it has not been investigated in dMMR GOAs whether the MSI hypermutator phenotype remains active during progression, how this impacts ITH and phylogenetic trees, and whether subclonal driver mutations evolve. Our previous work in kidney cancer for example showed that most driver mutations are located in subclones^12^. Subclonal driver mutations are suboptimal as therapeutic targets as co-existing wild type subclones remain untargeted^13^. They furthermore hinder effective biomarker development as the analysis of single tumour regions incompletely profiles the genomic landscape of the entire tumour. Large scale sequencing analyses of MSI GOAs identified *TP53, RNF43, ARID1A, PIK3CA, KRAS* and *PTEN*, as the most frequently altered driver genes. Mutations in antigen presentation (MHC class I, *B2M*)^2^ and interferon signalling pathway (*JAK1*/*2*)^*14,15*^ genes also frequently occur in MSI tumours and they have been suggested to enable immune evasion^2^. However whether they are truncal or subclonal within individual tumours has not been investigated.

Multi-region exome sequencing (MSeq) has been used to investigate cancer evolution by comparing mutational profiles from spatially separated tumour regions. MSeq found that mutations often appear to be present in all cancer cells (i.e. clonal) in a single tumour region even if they are absent from other regions of the same tumour^12,16^. Spatial constraints in solid tumours that preclude intermixing of evolving subclones likely explains this ‘illusion of clonality’ phenomenon when tumour heterogeneity is only investigated in a single sample per tumour^17,18^. Thus, MSeq can provide insights beyond single sample analysis. We applied MSeq to four surgically resected GOAs showing dMMR on immunohistochemistry and combined this with subclonality analysis of single tumour biopsies from each of 64 MSI GOAs sequenced by The Cancer Genome Atlas (TCGA)^2^ to assess ITH and the evolution of these tumours.

## RESULTS

Seven primary tumour regions from each of four GOAs (Fig. 1A) were subjected to MSeq with a minimum target depth of 200x (see Supplementary Table 1 for sequencing depths and mutation calls). Two lymph node metastases were included from each of two cases. TNM stage was assessed but no other clinical information was available as the samples had to be fully anonymised to comply with local ethics and research legislation. Absence of MLH1 and PMS2 staining and positive staining for MSH2 and MSH6 (Fig. 1B), indicated *MLH1* deficiency^19^. No known Lynch syndrome mutations in *MLH1, MSH2/6* or *PMS2* were identified in DNA from non-malignant tissue, confirming that these were sporadic rather than Lynch syndrome-associated dMMR tumours.

**Fig. 1.**
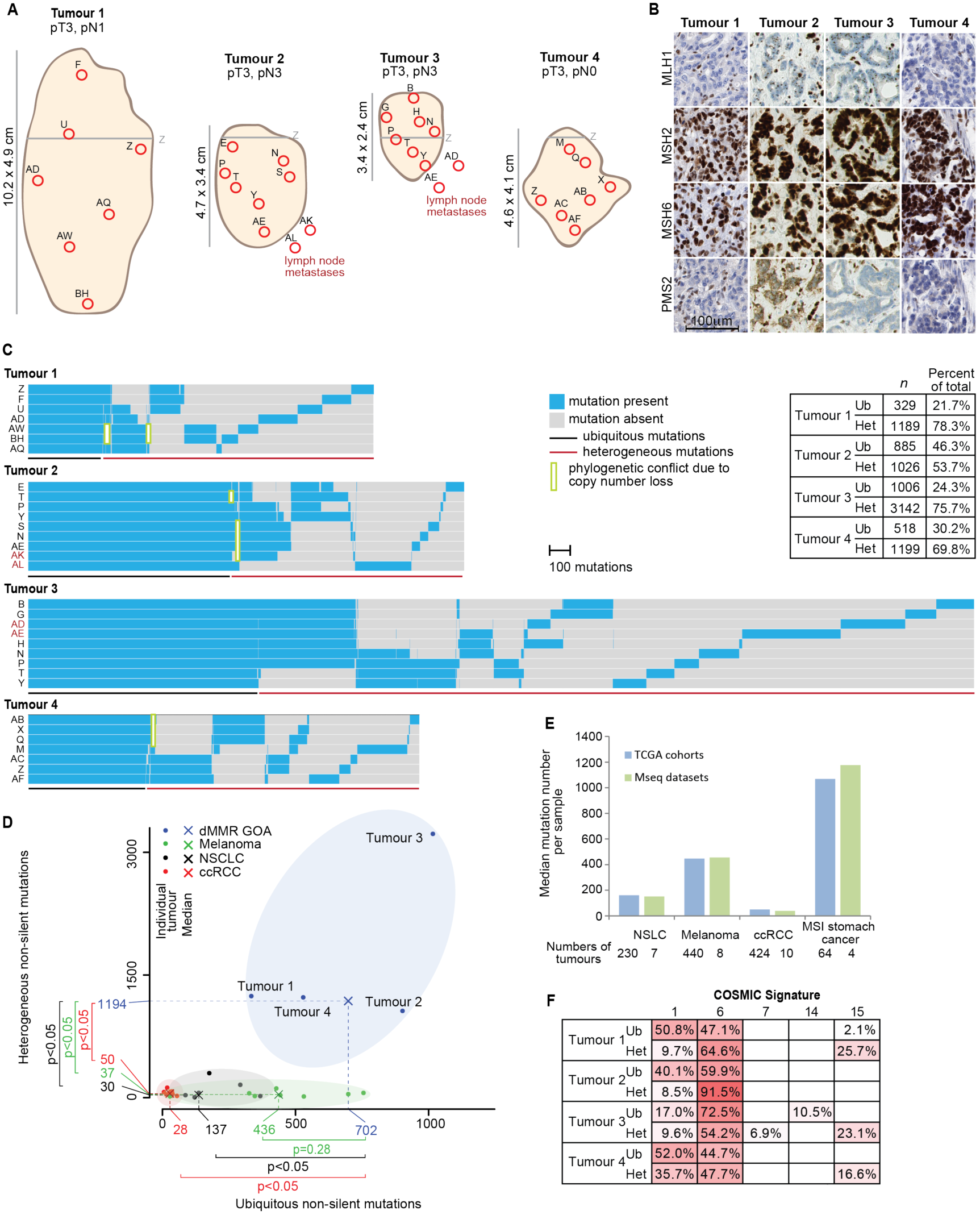
Intratumour heterogeneity of somatic mutations. **A:** Tumour size, location, TNM-stage and regions selected for sequencing. The grey line labelled (Z) marks the gastro-oesophageal junction. **B:** Immunohistochemical staining of MLH1, MSH2, MSH6 and PMS2. **C:** Heat maps showing the presence (blue) or absence (grey) of non-silent somatic mutations that were identified by MSeq across tumour regions. The table shows the number of heterogeneous (Het) and ubiquitous (Ub) mutations identified in each tumour and their percentage of the total non-silent mutation count of the tumour. **D:** Comparison of ubiquitous and heterogeneous mutation counts across four different tumour types analysed by MSeq (dMMR GOA: mismatch repair deficient gastro-oesophageal adenocarcinoma, Melanoma, NSCLC: non-small cell lung cancer, ccRCC: clear cell renal cell carcinoma). The Mann-Whitney test was used to assess significant differences in mutation loads between dMMR GOA and other tumour types. **E:** Median mutation loads of individual regions from MSeq datasets compared to the median single sample mutation loads from the Cancer Genome Atlas KIRC, SKCM, STAD and LUAD cohorts. **F:** COSMIC mutational signature analysis of ubiquitous (Ub) and heterogeneous (Het) mutations in four dMMR GOAs. Non-silent and synonymous mutations were included in the analysis and only signatures which contributed to ≥5% of mutations in at least one sample are shown.

### Mutational intratumour heterogeneity

1518 to 4148 (median: 1814) non-silent mutations were identified per case (Fig. 1C). The high mutation burden and the large fraction of indels among these mutations (20-34%) were consistent with a MSI phenotype^2^. The number of ubiquitous non-silent mutations mutations that were detected across all sequenced regions per tumour ranged from 329 to 1006 (median: 702). This exceeded the number of ubiquitous non-silent mutations reported for clear cell renal cell carcinomas (median: 28)^12^ and even for lung cancers (median: 137)^16^ and melanomas (median: 436)^20^, which are among the most highly mutated cancer types^21^ (Fig. 1D). The difference was significant between dMMR GOA and lung and renal cancers but not for melanomas. MSeq-identified ubiquitous mutations are likely to define the mutations that were present in the founding cell of each tumour before diversification into subclones occurred^12^. These high numbers hence reveal that the dMMR-phenotype was likely acquired in the pre-cancerous cell lineage considerably earlier (i.e. the time required for 329-1006 mutations to occur) than malignant transformation of the founding cell. Malignant transformation early after the acquisition of the dMMR-phenotype which is then followed by selective sweeps that eradicated early heterogeneous subclones could be an alternative explanation. Yet, it appears unlikely that such a process would have left no trace of the early subclones in any of these tumours. A median of 1194 non-silent mutations were not detectable across all regions per case and hence heterogeneous. This significantly exceeded the heterogeneous mutation burden detected by MSeq in ccRCC^12^ by 24-fold, in lung cancer^16^ by 39-fold, and in melanoma^20^ by 32-fold (Fig. 1D). Importantly, the median mutation load per analysed region in these MSeq series was similar to that reported by the TCGA for the respective cancer type (Fig. 1E), suggesting that these small series are reasonably representative of each tumour type. Thus, dMMR tumours are characterized by extreme ITH compared to the other three cancer types.

High mutation and neoantigen loads are associated with immunotherapy benefit. Recent data suggested more specifically that a high burden of clonal mutations/neoantigens is important for immunotherapy success ^22,23^. Applying the NetMHC algorithm predicted 1120 to 3052 strong class I MHC binding neoantigens per tumour (Supplementary Fig. 1). Between 215 and 926 of these were clonal neoantigens. This is higher than clonal neoantigen loads reported for most lung cancers or melanomas^24^. It is conceivable that this high clonal neoantigen burden explains the immunotherapy sensitivity of dMMR tumours^22^.

### Mutational signatures reveal mutational processes driving evolution

We next investigated mutational signatures by counting the number of all possible base substitution in their trinucleotide contexts (Supplementary Fig. 2) and then assigning these to 30 COSMIC mutational signatures^21^ (Fig. 1F). The COSMIC mutational signatures 6 and 15 are characteristic for MSI cancers and these were abundant among ubiquitous *and* heterogeneous mutations. Signature 1 mutations reflect the spontaneous deamination of methylated cytosine, a mutational process active in most normal tissues. Signature 1 was detected in 17-52% (219-449 mutations in absolute number) of ubiquitous mutations. A fraction of these were likely acquired in the normal cells over the lifetime of these patients. However, based on the estimated mutation acquisition rate in normal gastro-oesophageal epithelium, only 0.5-1 signature 1 mutations would be expected to accumulate per lifeyear^25-27^. It is hence likely that the dMMR phenotype also contributes to the generation of signature 1 mutations. This is further supported by 9-10% of the subclonal mutations in Tumours 1-3 and 36% in Tumour 4 showing signature 1 and consistent with a recently suggested role of the MMR system in the repair of deamination defects^28^. No other mutational signatures contributed substantially to the heterogeneous mutations, confirming that the MSI-phenotype remains active during cancer progression and is the primary mechanism generating these large numbers of subclonal mutations.

### The evolution of copy number aberration landscapes

DNA copy number aberration (CNA) profiles revealed near-diploid profiles across all regions of Tumour 2 and Tumour 3 (Fig. 2A and Supplementary Fig. 3). Tumour 4 showed highly aberrant and near-tetraploid profiles in all regions. A high number of mutations were present on all copies of the major allele of most gained chromosomes (Fig. 2B), indicating that whole genome duplication and chromosomal instability (CIN) had occurred late on the trunk of the phylogenetic tree in Tumour 4. CIN was confirmed by the wGII index that measures the proportion of all chromsomes with copy number states that differ from the ploidy of a sample and where values above 0.2 support the presence of CIN^29^ (Fig. 2A). Near-diploid *and* near-triploid CNA profiles were found in distinct regions of Tumour 1. Together with an increase in wGII from values ∼0.2 in the near-diploid regions to over 0.5 in near-triploid regions and the occurrence of new CNAs in individual tumour regions, this revealed the acquisition of subclonal CIN during cancer progression. Importantly, all four analysed lymph node metastases were near-diploid with wGII values ≤0.2, demonstrating that CIN, which has been associated with tumour aggressiveness in several cancer types including GOA^30^, is not required for metastasis formation.

**Fig. 2.**
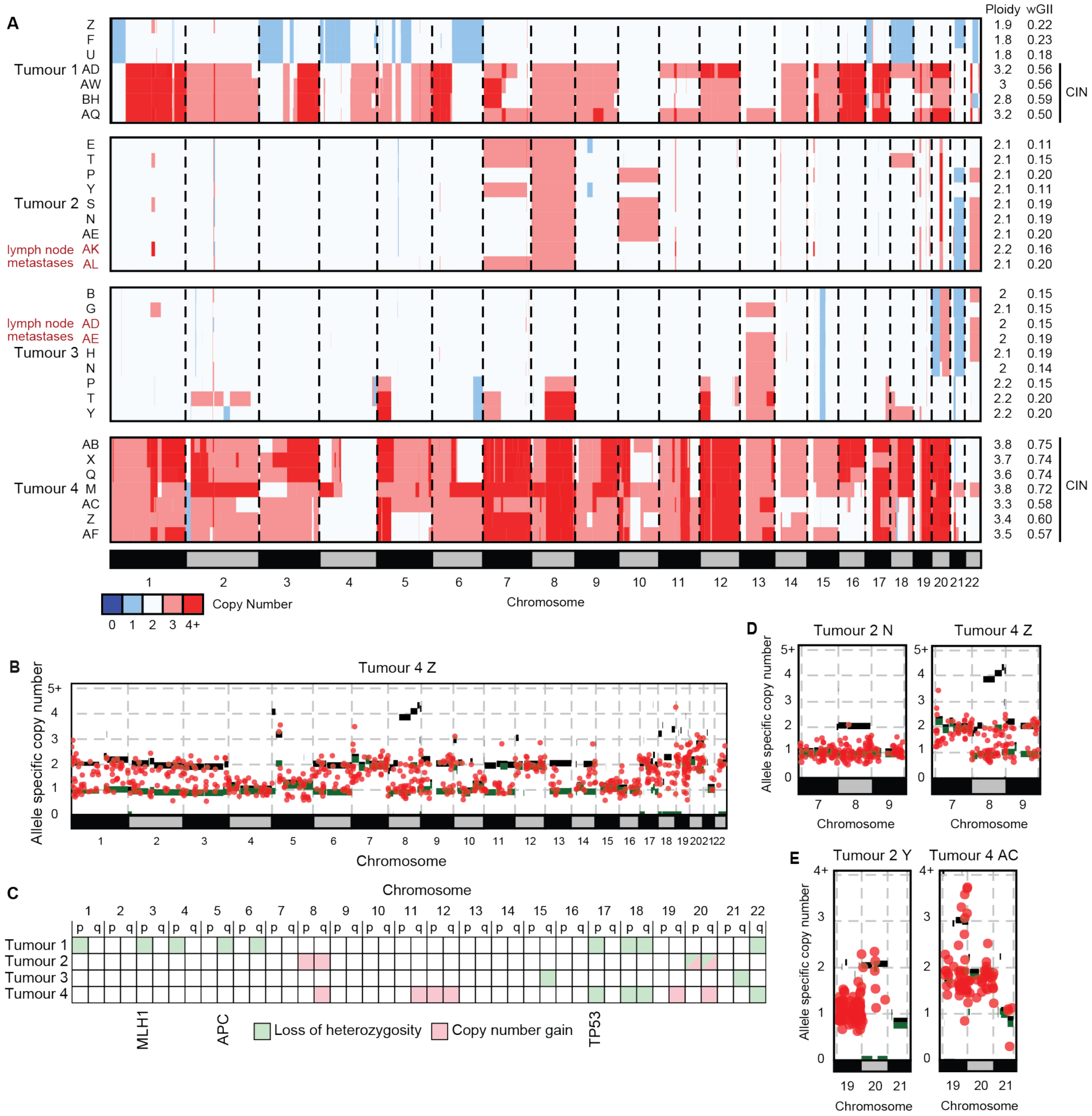
Intratumour heterogeneity of DNA copy number aberration. **A**: Genome-wide DNA copy number profiles of each tumour region. Profiles showing chromosomal instability (CIN) are labelled with a black bar on the right. **B**: Example of an allele specific DNA copy number profile and superimposed copy numbers of somatic mutations from Tumour 4. This allows timing of CIN/genome duplication, demonstrating late acquisition, as large numbers of mutations are located on the major alleles for most gained chromosomes. **C**: Ubiquitous loss of heterozygosity (LOH) or copy number gains identified in each of the four tumours. Tumour suppressor genes commonly mutated in dMMR GOAs and which are located on chromosomes showing ubiquitous LOH are labelled. **D**: Examples of the allele specific copy number and copy number of corresponding somatic mutations for Chr8 and **E**: for Chr20 which showed recurrent ubiquitous gains in our series.

We next investigated which specific CNAs were ubiquitous/clonal and had hence occurred early in the evolution of these dMMR tumours (Fig. 2C and Supplementary Fig. 3). Ubiquitous Chr17p, Chr18 and Chr22 loss of heterozygosity (LOH) events were each present in two tumours. Ubiquitous LOH events of Chr3p, Chr5q and Chr17p encompassed tumour suppressor genes which are recurrently mutated in dMMR GOAs^2^ (*MLH1, APC* and *TP53*, respectively). Among the small number of ubiquitous gains, only Chr8q and Chr20q gains were identified in more than one tumour. To further time the acquisition of these recurrent truncal CNAs in this small series, we mapped ubiquitous mutations onto the allele-specific CNA profiles. Copy number gains that occurred early can be identified if the majority of mutations in that region have a copy number which is lower than that of the gained allele. The Chr8 gain in Tumour 2 and the Chr8q gain in Tumour 4 (Fig. 2D), but not Chr20 gains (Fig. 2E), showed a near complete absence of mutations and were hence acquired on the phylogenetic trunk before or soon after the MSI-phenotype. Thus, Chr8q gains, which are the commonest CNAs in MSI GOAs^2^, can be among the earliest genetic aberrations in these tumours.

### Reconstruction of tumour phylogenetic relationships

We next deconvoluted the subclonal composition of individual tumour regions and reconstructed the phylogenetic tree for each tumour (Fig. 3). Similar to MSeq analyses of other tumour types^12,16,20^, this revealed branched evolution. Comparison of the phylogenetic trees with the mutation heatmaps showed some phylogenetic conflicts. Inspection of the copy number status of the mutated DNA positions showed that most conflicts could be explained by losses of chromosome copies in individual regions (marked in green in Fig. 1C and Supplementary Fig. 4). Thus, subclones can lose a small proportion of mutations during cancer evolution.

**Fig. 3.**
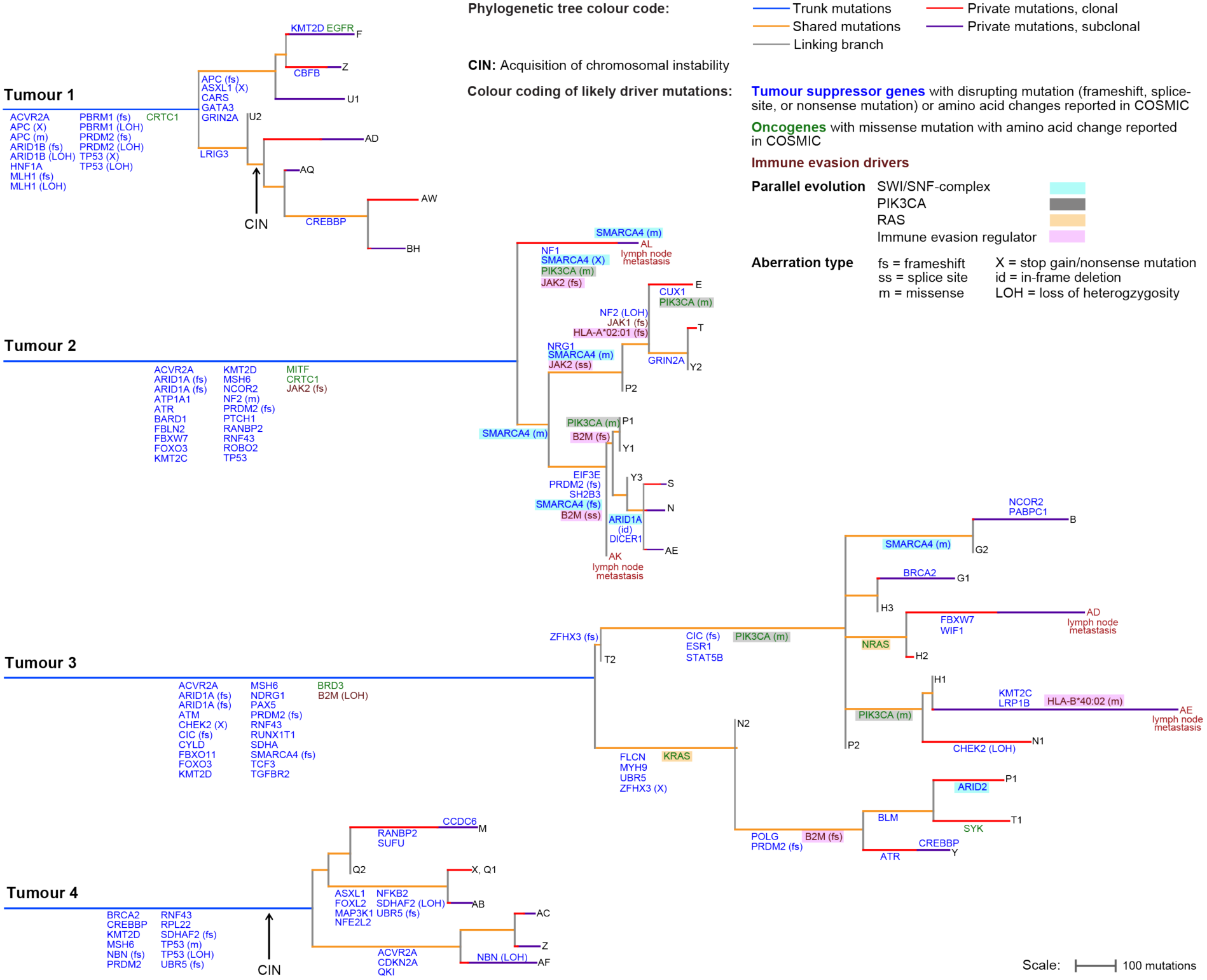
Tumour phylogenetic trees. Trees were reconstructed from non-silent *and* synonymous mutations and trunk and branch lengths are proportional to the number of mutations acquired. Trees are rooted at the germline DNA sequence, determined by exome sequencing of DNA from tumour adjacent normal tissue. Subclones that define the tips of the tree are labelled with the tumour region in which they were identified. Numbers were added where several subclones were identified by the phylogenetic deconvolution algorithm within a tumour region, with 1 defining the largest intra-regional subclone and 2, 3 increasingly smaller subclones. Private mutations were furthermore split into those that were clonal in the analysed region (present in >0.7 of the cancer cell fraction in that region) and those that were subclonal (present in ≤0.7 of the cancer cell fraction). Likely driver mutations and relevant loss of heterozygosity (LOH) events were mapped onto the branch of the trees where they likely occurred. Genes affected by more than one genetic aberration within a tumour are labelled with the genetic aberration type that occurred. Arrows labelled ‘CIN’ indicate the likely onset of chromosomal instability.

Phylogenetically closely related clones were located in proximity to each other within tumours (Supplementary Fig. 5), indicating that cell motility is limited and that these solid tumours evolve in a spatially ordered fashion. Importantly, each of the two lymph node metastases analysed in Tumours 2 and 3 had evolved from distinct subclones rather than being seeded by the same subclone or sequentially from one node to the other (Fig. 3). Dissemination hence propagated a proportion of the subclonal diversity from the primary tumour to metastatic sites. In addition, subclonal mutations, defined as private mutations estimated to be present in ≤70% of the cancer cells of a sample, were detectable *within* three metastatic sites with good cancer cell content (37-82% cancer cell content, Supplementary Table 2) but not in the metastasis with low cancer cell content (15% in sample AK, Tumour 2) which likely precluded the detection of subclonal mutations. The subclonal mutations *within* lymph nodes were again predominated by the MSI-specific mutational signatures 6 and 15 (Supplementary Table 3). Thus, the dMMR phenotype continues to generate ITH in metastases.

### Identification of truncal driver aberrations

We next assessed the evolution of putative driver mutations and of corresponding LOH events of tumour suppressor genes and mapped them onto the phylogenetic trees (Fig. 3 and Supplementary Table 4 for details of putative driver mutations). A frameshift mutation and LOH of *MLH1* had occurred on the trunk of Tumour 1, consistent with biallelic *MLH1* loss. No genetic aberrations of *MLH1* were detectable in Tumours 2-4 even though immunohistochemical analysis had demonstrated *MLH1* deficiency (Fig. 1B). Hypermethylation of *MLH1* is hence the likely cause for dMMR in these cases. Tumours 2-4 furthermore harboured a truncal frameshift mutation in *MSH6* but no LOH of *MSH6* was identified and the functional relevance of monoallelic *MSH6* loss is unclear. Mutations in the histone methyltransferase and tumour suppressor gene *PRDM2*, one in combination with LOH of the second allele were also truncal in all 4 cases and truncal frameshift mutations of the TGFβ signalling regulator *ACVR2A* were detected in three cancers.

One tumour showed a disrupting mutation and LOH of *ARID1B* and two tumours each harboured two mutations in *ARID1A* on the trunk, which are all members of the SWI/SNF chromatin-modifying complex. We could not formally demonstrate that the two mutations affected both alleles of the *ARID1A* tumour suppressor gene but biallelic inactivation is likely as all mutations were disrupting in nature, suggesting evolutionary selection for inactivating events. A frameshift mutation and LOH of *PBRM1*, a further SWI/SNF-complex member, co-occurred with biallelic *ARID1B* loss on the trunk of Tumour 1. This further emphasizes an important role for SWI/SNF-complex aberrations in dMMR GOA development.

Truncal mutations in *TP53* were found in three tumours. Two of these (Tumours 1 and 4) also showed LOH, leading to biallelic *TP53* inactivation. These specific cancers had undergone genome duplication and acquired CIN, consistent with a permissive role of *TP53* loss for CIN^31^. Moreover, both showed truncal Chr18q loss which promotes CIN in colorectal cancer^32^. *TP53* inactivation and Chr18q loss may hence predispose tumours to subsequently evolve CIN. Frameshift mutations of *RNF43*, a negative regulator of the APC/β-catenin-pathway that frequently acquires heterozygous mutations in MSI tumours^33^, were present in three tumours. The tumour without an *RNF43* mutation harboured two truncal mutations in the *APC* tumour suppressor gene as an alternative mechanism of β-catenin activation. Together, aberrations in *TP53*, the SWI/SNF-complex, *PRDM2*, dMMR-, APC/β-catenin signalling- and TGFβ signalling-genes each occurred on the phylogenetic trunks of two or more tumours.

### Identification of parallel evolution

Assessing heterogeneous driver mutations revealed striking examples of parallel evolution in two tumours, a strong signal that these evolved through Darwinian selection^17^: Tumour 2 acquired five subclonal mutations in *SMARCA4*, encoding a catalytic subunit of the SWI/SNF-complex. These had occurred in addition to two truncal mutations (M274fs, K1071fs) in the *ARID1A* SWI/SNF-complex member. A third *ARID1A* mutation was subclonal and affected recurrently mutated amino acids (AA163-164 deletion) proximally to the truncal frameshift mutations. This may be functionally relevant if *ARID1A* had retained some residual function despite the more distal mutations. Together, parallel evolution of five *SMARCA4* mutations in this tumour with truncal *ARID1A* mutations suggests that SWI/SNF-complex aberrations are not only important for carcinogenesis but that progressive inactivation of the SWI/SNF-complex may contribute to cancer progression.

A *PIK3CA* hotspot mutation (H1047R) was detected in P1 and Y1 but also in the distantly related subclone AL in Tumour 2. Copy number changes that could explain a loss of this mutation in subclones with wild-type *PIK3CA* were absent (Supplementary Fig. 3). The most parsimonious explanation for this phylogenetic conflict is that the same mutation independently evolved twice, once in AL and once in the ancestor cell of P1 and Y1. With one further *PIK3CA* hotspot mutation in region E (Y1021C), this identified three *PIK3CA* parallel evolution events in Tumour 2.

Mutations in the SWI/SNF-complex members *SMARCA4* and *ARID1A* were present on the trunk of Tumour 3. Additional SWI/SNF mutations, one in *ARID2* and one in *SMARCA4*, evolved in subclones, the latter potentially complementing monoallelic *SMARCA4* loss on the trunk to biallelic inactivation in the subclone. Further parallel evolution was apparent in Tumour 3 based on the acquisition of *KRAS* (G13D) and *NRAS* (G12C) oncogenic mutations in distinct subclones. Two hotspot *PIK3CA* mutations (E418K, Y1021H) sequentially occurred in one clade of Tumour 3.

The tumour suppressor gene *PRDM2* harboured frameshift mutations on the trunks of Tumours 2 and 3 and a second frameshift mutation was acquired in subclones of each tumour, potentially leading to biallelic inactivation. Subclonal inactivating mutations of the cell cycle regulator and DNA damage repair genes *CHEK2, ATR* and *BLM* occurred in Tumour 3. Together with truncal LOH of *CHEK2*, both alleles of this gene were inactivated. Heterozygous mutations of *BLM* and *ATR* may be functionally relevant as both genes show haploinsufficiency^34,35^.

Given the very high burden of substitutions and indels caused by MMR deficiency, it is likely that several of the mutations we classified as likely drivers are passengers without a significant fitness effect. However, parallel evolution and the strong functional evidence for driver status of the identified *KRAS, NRAS* and *PIK3CA* mutations and of inactivating mutations in SWI/SNF-complex members in cancer^36^ support the functional relevance of these specific aberrations.

### The evolution of immune evasion drivers

Tumour 2 harboured a truncal *JAK2* frameshift mutation. In addition, a subclonal *JAK2* splice-site mutation evolved in one clade of this tumour and a further frameshift mutation in region AE. Another subclone had acquired a heterozygous *JAK1* frameshift mutation but no evidence for biallelic inactivation was found. A subclonal frameshift mutation was present in *HLA-A*\**02:01* (Supplementary Table 5). Assessing the neoantigens binding to this HLA allotype revealed that this would lead to a 12% reduction in the number of neoantigens presented by the affected subclones (Supplementary Fig. 6). One clade in Tumour 2 furthermore acquired two disrupting mutations in *B2M*. Both mutations were located in close proximity and, when inspecting short read sequencing data, these were not located on the same allele but conferred biallelic inactivation which abrogates MHC Class I antigen presentation (Supplementary Fig. 7).

LOH of *B2M* was present on the trunk in Tumour 3 and a *B2M* frameshift mutation was acquired in a subclone, also establishing biallelic *B2M* loss. Although several primary tumour regions in Tumours 2 and 3 showed biallelic *B2M* inactivation this was not propagated to any of the four lymph node metastases (Fig.3). The lymph node metastasis AE in Tumour 3 acquired a missense mutation in *HLA-B*\**40:02* (Supplementary Table 5). Compared to the frameshift mutation in *HLA-A*\**02:01*, the functional impact of this mutation is unknown. If this *HLA-B*\**40:02* mutation compromised antigen presentation, 12% of neoantigens could no longer be presented. In contrast to lung cancers which are frequently chromosomally unstable and acquire subclonal LOH of *HLA* genes as immune evasion mechanisms^37^, no such LOH events were identified (Supplementary Table 5).

To investigate why immune evasion drivers only evolved in two of the four tumours, we assessed cytotoxic CD8+ T-cell infiltrates by immunostaining. The two tumours with evidence of immune evasion events, which also had the highest truncal and subclonal mutation burdens, showed higher T-cell infiltrates compared to the other two tumours (Fig. 4). dMMR GOAs with high immunogenicity and T-cells infiltrates may hence be particularly prone to subclonal immunoediting.

**Fig. 4.**
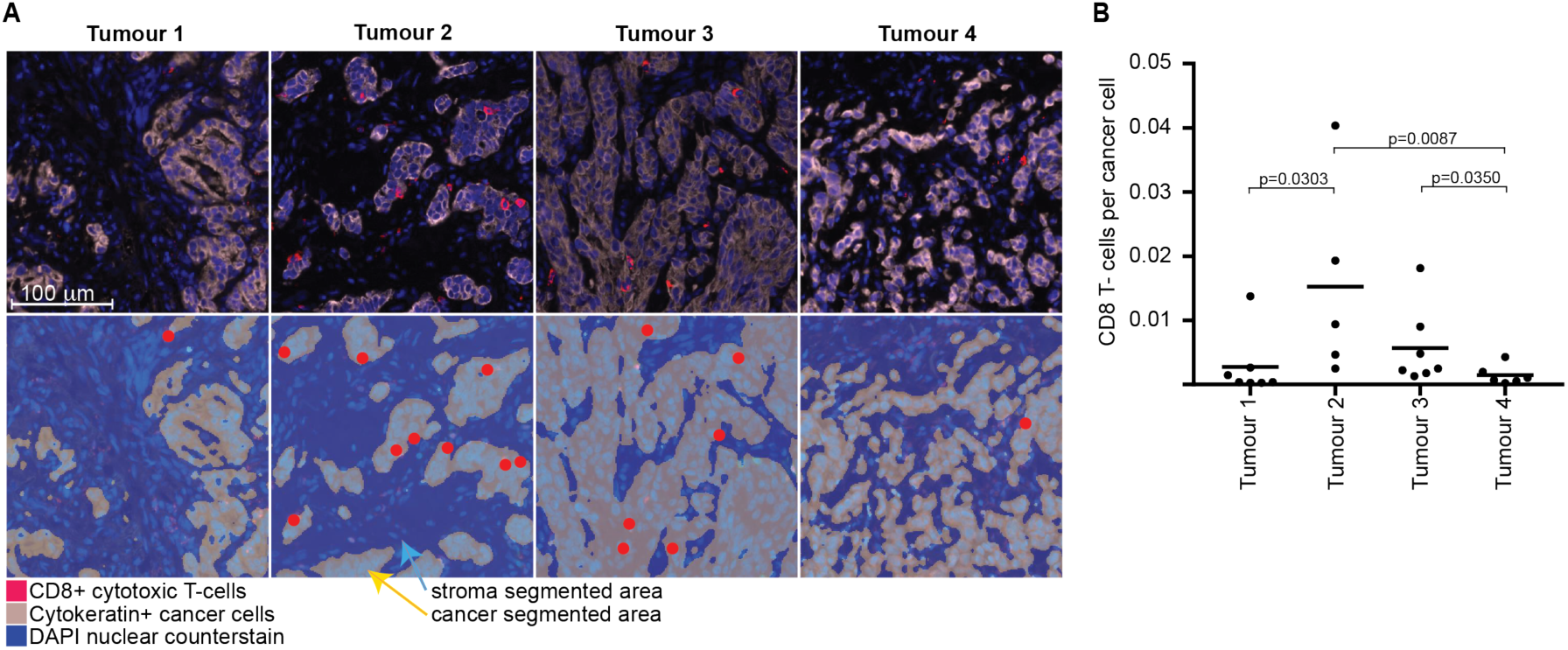
CD8+ T-cells infiltrates in dMMR GOAs. **A**. Representative images of CD8+ T-cell infiltrates in Tumours 1-4. Upper row: fluorescent composite IHC image showing cancer cells and CD8+ cytotoxic T-cells. Bottom row: Segmentation of cancer areas (grey, yellow arrow) and stroma (blue, light blue arrow) allowed to only count the highlighted CD8+ T-cell in cancer areas. **B**. The ratio of CD8+ T-cells divided by the number of cancer cells (Cytokeratin positive cells) for all regions of Tumours 1-4. Black bar: median. p-values (Spearman rank test) are shown for significant differences.

### Non-synonymous to synonymous mutation ratio evolution

The ratio of non-synonymous mutations to synonymous mutations (dN/dS ratio) has been used to estimate positive and negative selection in cancer^38^. dMMR tumours have high clonal but also subclonal mutation burdens and we reasoned that this may enable applying these ratios to evaluate how selection changes from truncal mutations to subclones. dN/dS ratios were close to 1 for the truncal mutations of all cases (0.95-1.06), indicating that the majority of mutations are neither under positive nor under negative selection. However, the dN/dS ratios increased to 1.16 in Tumour 1 and 1.31 in Tumour 2 for private mutations, indicating positive selection (Fig. 5 and Supplementary Table 6). Together with the identification of parallel evolution in Tumours 2 and 3, this strongly suggests that these tumours are under selection pressure and that adaptive mutations continue to evolve.

**Fig. 5.**
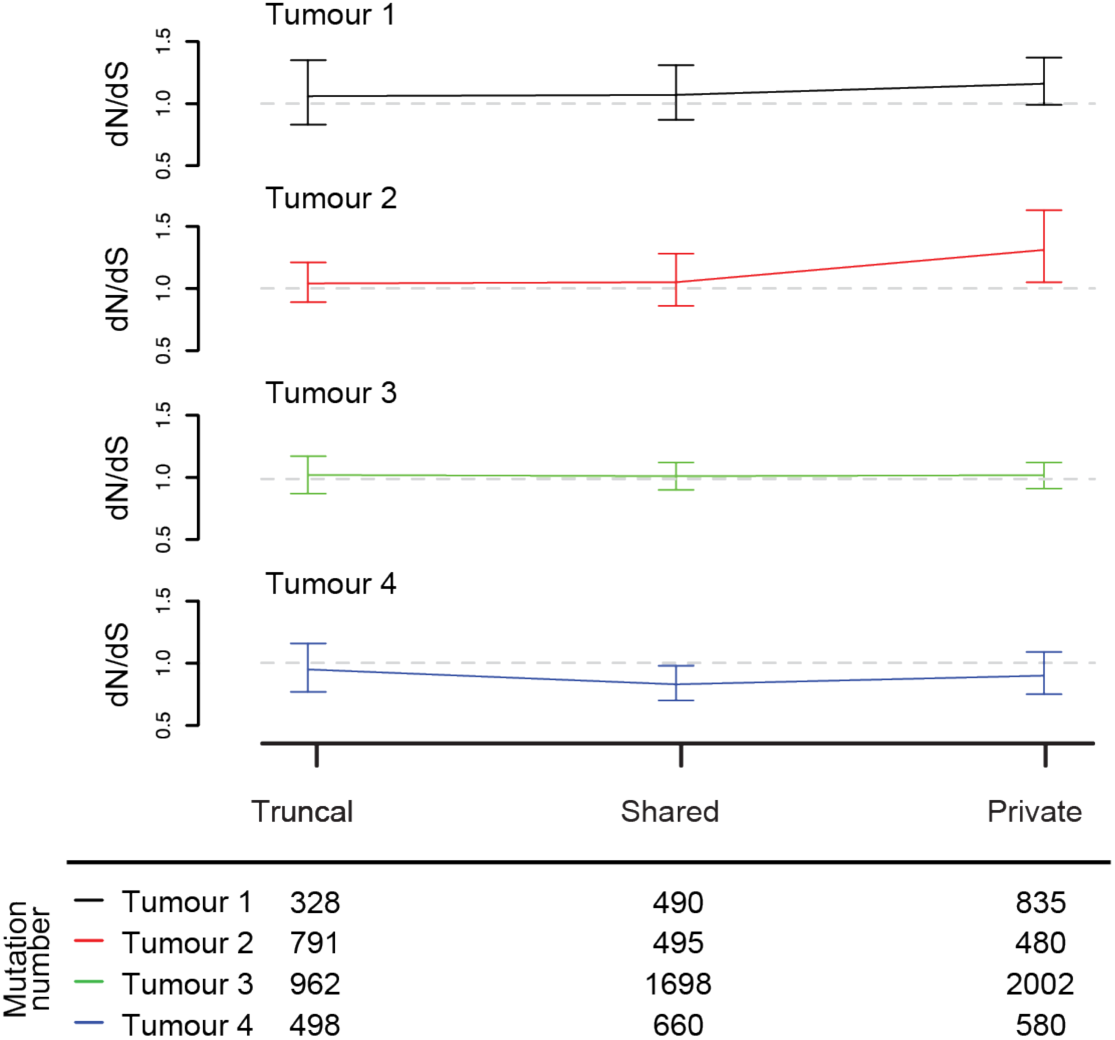
Non-synonymous to synonymous (dN/dS) mutation ratios for ubiquitous, shared and private mutations. dN/dS ratios adjusted for common mutation biases. Error bars show 95% CI’s. The total number of synonymous and non-synonymous mutations available in each category for the analysis are shown beneath the plot.

### Comparison of multi-region vs single-regions heterogeneity analysis

Our next aim was to gain further insights into the evolution of dMMR GOAs by deconvolution of clonal and subclonal mutations in single samples from the TCGA GOA dataset^2^. However, we first used our MSeq dataset to assess which information can be robustly generated by single samples deconvolution and which ones are more likely to be gained by MSeq. The total mutation load in a single sample exceeded the MSeq-determined ubiquitous/truncal mutation load by an average of 73% across the four tumours (Fig. 6A). Following bioinformatic deconvolution of mutations in each individual region into clonal and subclonal, the average clonal mutation burden determined in single samples still exceeded the number of mutations identified as ubiquitous by MSeq by 34%. Importantly, the number of mutations identified as clonal in a single region varied highly between samples from the same tumour. For example, clonal mutation numbers in Tumour 1 were only overestimated by 15% in region U but by 70% in region BH compared to MSeq-data. This could not be attributed to different cancer cell contents between regions as no correlation was observed (Supplementary Fig. 8).

**Fig. 6.**
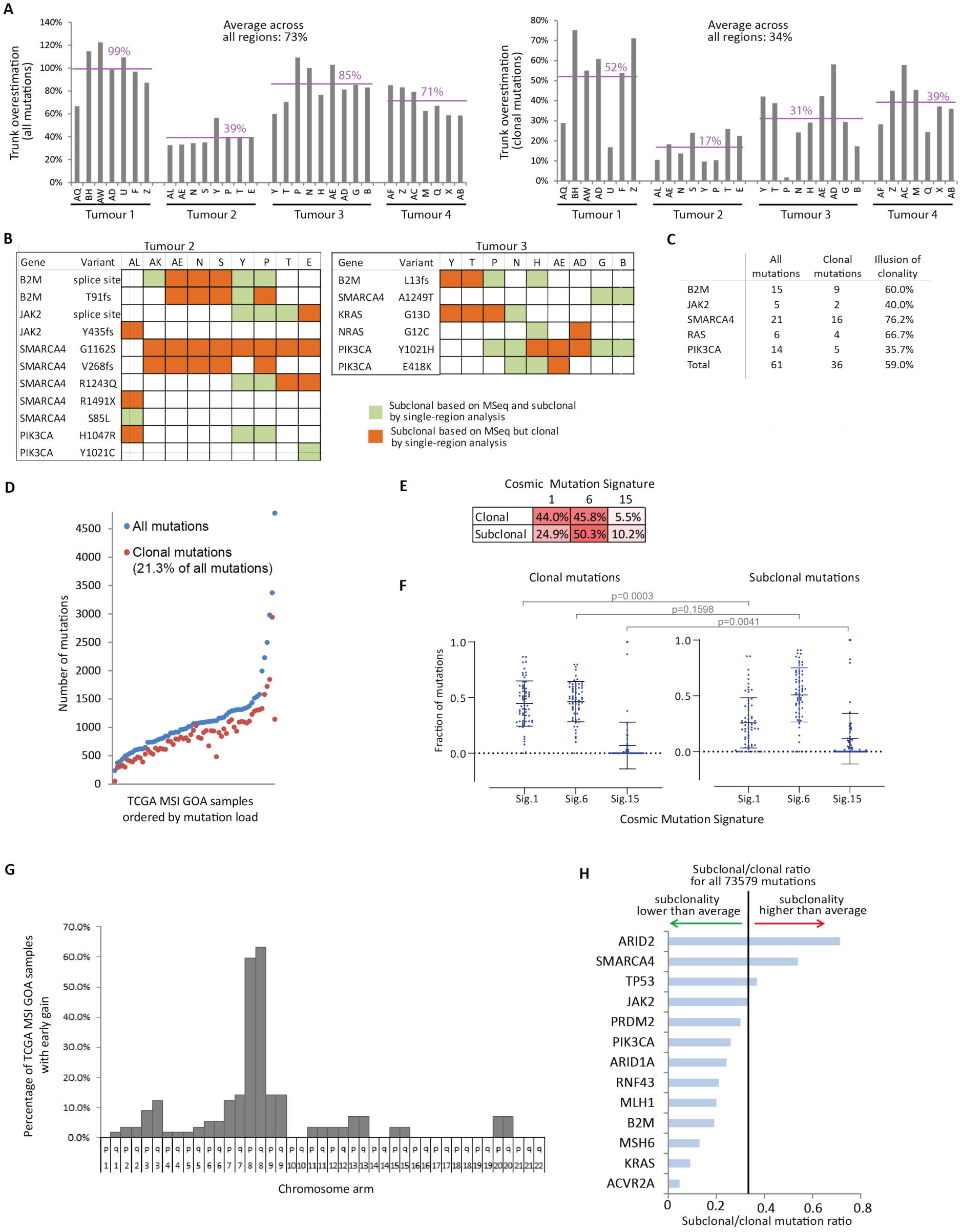
Clonal and subclonal mutation analysis. **A:** Comparison of the total non-silent mutation load per region and of the clonal mutation load per region (defined as present in a cancer cell fraction (CCF) of 0.7 or above) against the number of ubiquitous mutations that have been identified by MSeq. The percent difference to the ubiquitous mutation load is shown. **B**. Subclonal driver gene mutations assessed by MSeq in Tumour 2 and 3. In green mutations that single region analysis picks up as subclonal and in orange mutations that would have been falsely assigned as clonal by single region analysis. **C**. Illusion of clonality (in percent) for driver gene mutations. **D**. Mutation load of TCGA MSI GOA samples, number of all mutations in blue and of clonal mutations in red. **E**. Mean percentage of clonal and subclonal mutational signatures found in TCGA MSI GOA samples. **F**. Subclonal and clonal mutational signatures for 64 TCGA MSI GOAs. Means and standard deviation are shown and p-values have been calculated with a Mann-Whitney test. **G**. Percentage of the 64 TCGA samples that gained the indicated chromosome arm early, i.e. before a high number of mutations was acquired through MSI. **H**. Subclonal to clonal mutation ratio for driver gene mutations. The black line shows the average ratio across all somatic mutations.

We furthermore assessed whether the parallel evolution mutations, that have a high probability of being actual drivers and were found to be subclonal by MSeq analysis, could also have been accurately identified as subclonal by single region analysis. Only 40% of *B2M* mutations that were subclonal based on MSeq were accurately identified as subclonal in individual regions whereas 60% appeared clonal (Fig. 6B-C). This illusion of clonality in single sample analysis also affected 40% of *JAK2* mutations, 76.2% of *SMARCA4* mutations, 66.7% of *RAS* mutations and 35.7% of *PIK3CA* mutations. Overall, 59.0% of these likely driver mutations appeared clonal in single region analysis despite clear suclonal status when considering the MSeq data. This supports the conclusion from MSeq studies in other tumour types, that single region analysis often overestimates the clonal dominance of driver mutations^12,16^.

Having established these limitations of single-region heterogeneity analyses, we analysed the 64 MSI GOAs cancers from TCGA. All samples harboured subclonal mutations and a median of 21.3% of mutations were subclonal (Fig. 6D). Mutational signature analysis showed that clonal and subclonal mutations were dominated by MSI-specific mutational signatures 6 and 15 (Fig. 6E-F), confirming our MSeq results. 44.0% of clonal mutations displayed signature 1 and although this significantly decreased among subclonal mutations, it remained the second most abundant mutation signature. Together with a significant increase in signature 15 among subclonal mutations, this supports the change in mutational processes between early progression and subclonal diversification as seen in the MSeq dataset.

Timing of copy number changes in the TCGA dataset furthermore supported that chromosome 8 gains had been acquired before or early after the MSI-phenotype in ∼60% of cases (Fig. 6G and Supplementary Fig. 9). We finally investigated the clonality of mutations in driver genes that were predominantly clonal or subclonal by MSeq also in the TCGA samples. The highest frequency of subclonal mutations was found in *SMARCA4* and *ARID2* whereas *ACVR2A* was almost always clonal in TCGA data (Fig. 6H), consistent with MSeq data where these occurred late and early, respectively. Mutations in the remaining driver genes were predominantly clonal in TCGA data, but this is likely limited by the overestimation of clonal status in single region analysis. Of note, two independent disrupting mutations in *ARID1A* were found to be clonal in each of 16 out of 64 tumours (25%) and only 4 tumours had one clonal and one subclonal inactivating event. Combined with our observation of truncal *ARID1A* alterations, this may support early biallelic inactivation in a fraction of these tumours.

### Mutational mechansism and their timing influence cancer phylogenetic tree morphologies

To investigate how mutational processes and their timing influence phylogenetic tree morphologies, we represented dMMR GOAs, melanomas^20^, lung^16^ and renal cancers^12^ as a single phylogenetic tree with a branching structure similar to those revealed by MSeq and by using the average number of ubiquitous and heterogeneous mutations (Fig. 1D) to scale trunk and branch sizes (Fig. 7). This revealed that dMMR leads to long trunks even exceeding the trunk size of typically carcinogen induced cancers (UV radiation in melanomas, cigarette smoke in lung cancer). Additionally the dMMR phenotype showed prominent branches whilst branch length in lung cancer and melanoma were similarly short as in renal cancers^12^, a consequence of the limited impact of the initiating carcinogens during cancer progression^16,20^. These associations show that mutation rates and their temporal activity are major factors determining phylogenetic tree shapes and sizes.

**Fig. 7.**
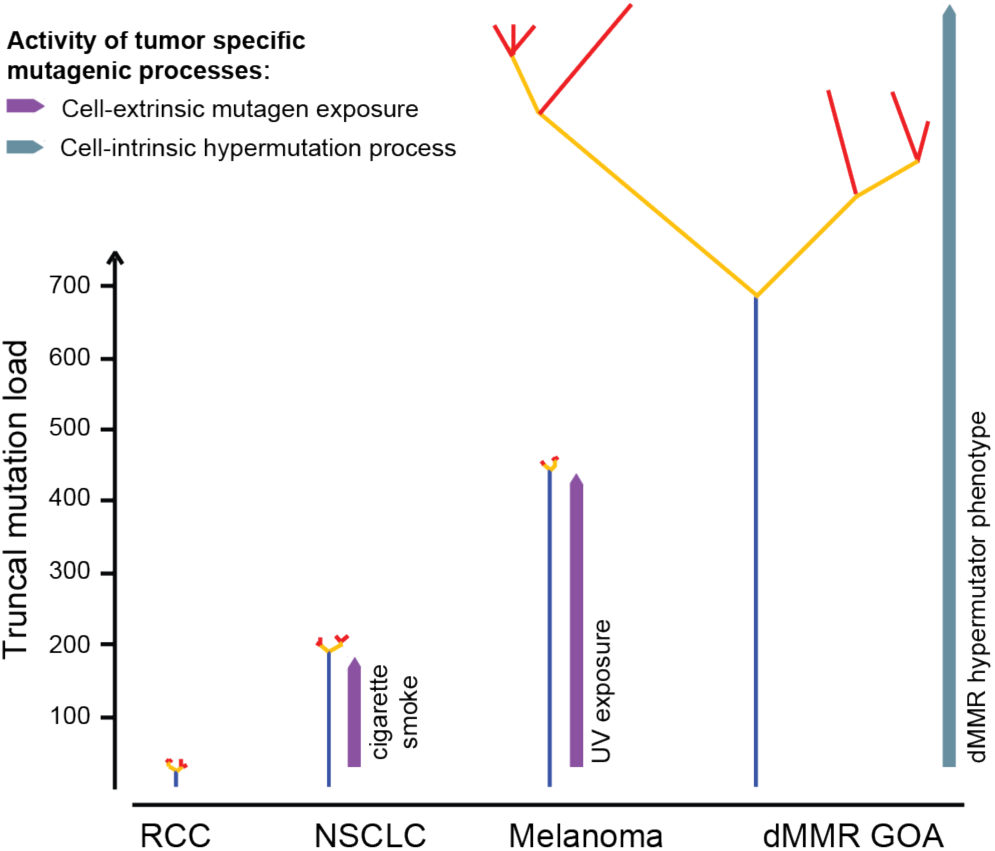
Comparison of phylogenetic tree morphologies across four cancer types analysed by MSeq. Schematics of branched phylogenetic trees drawn with similar branching structures to those directly observed in each of the four tumour types ^12,16,20^. Trees were scaled so that trunk and branch lengths are proportional to the average number of ubiquitous and heterogeneous non-silent mutation loads of each tumour type (Fig. 1C). Phylogenetic tree colour code: blue: truncal mutations, yellow: shared mutations, red: private mutations.

## Discussion

With recent success rates of cancer-immunotherapy, understanding the genetic landscapes of immunotherapy sensitive tumour types and how these influence treatment sensitivity are major needs. dMMR cancers are among the most sensitive solid tumours to checkpoint-inhibiting immunotherapies^7,8^ but their genetic evolution, clonal mutation burden and ITH remained unknown. Our series of four treatment-naïve dMMR GOAs revealed strikingly high clonal mutation burdens. This may explain the exquisite sensitivity of these cancers to immunotherapy as recent data showed that a high clonal mutation burden is a better predictor of immunotherapy success than the total mutation burden^22^. The presence of mutational ITH has furthermore been suggested to impair effective immunotherapy in lung cancer and other malignancies^22,23^. Extremely high numbers of heterogeneous mutations were found in all four dMMR GOAs and these significantly exceeded those in other cancer types analysed by MSeq. Although the analysed tumours were not treated with immunotherapy, these results and the overall high response rate of dMMR GOAs suggests that extreme ITH is unlikely to fundamentally preclude immunotherapy efficacy in tumours with abundant clonal mutations.

Our study also provided first insights into the clonal origin of lymph node metastatic disease in dMMR GOAs. Lymph nodes were seeded by distinct subclones in the primary tumours, propagating some of the heterogeneity from the primary tumour to metastatic sites. Subclonal mutation generation continued in metastases and similar heterogeneity as observed in primary tumours should therefore be expected in more advanced metastatic disease.

The mutation load of individual tumour regions exceeded the number of truncal mutations by 73%, and still by 34% following subclonal deconvolution. Studies investigating mutation burden as immunotherapy biomarkers may hence benefit from MSeq to robustly and accurately estimate truncal mutation loads. Subclonal immune evasion drivers were identified in two of four cases. Mutations in the *JAK1/2* and inactivation of *B2M* can confer resistance to checkpoint-inhibiting immunotherapy^15,39,40^. Although in MSI colorectal cancer it has been shown that most patients with *B2M* inactivation benefitted from immunotherapy^41^, our data suggests that *B2M* loss can be subclonal and is not necessarily propagated to metastasis. How subclonal immune evasion drivers and their localisation in the primary tumours or in metastases impairs immune checkpoint-inhibitor efficacy in dMMR GOAs should be investigated by MSeq in larger, immunotherapy treated cohorts.

Despite the selection pressure resulting from the high immunogenicity of dMMR tumours, we found no evidence of reversion of the hypermutator phenotype. Immune evasion mechanisms which can be readily accessed through single mutations, for example in *HLA* genes, or through biallelic *B2M* or *JAK* mutations may more effectively mitigate against this selection pressure than loss of the dMMR-phenotype which would still leave behind neoantigen-encoding mutations that have already been generated. Despite considerable mutation loads, cytotoxic T-cell infiltrates were low in two tumours and we could not identify immune evasion events that explain this. This warrants further investigation into immune escape mechanisms in dMMR GOAs.

Defining driver mutations which are commonly truncal is critical for precision cancer medicine approaches as targeting of subclonal driver mutations is likely futile^13^. Several tumour suppressor genes were inactivated by genetic alterations on the trunk in all four tumours. However, loss of function of tumour suppressor genes is usually not directly targetable. 2 of 4 dMMR GOAs harboured two inactivating mutations in *ARID1A*. In addition, 25% of MSI GOAs from the TCGA dataset showed two clonal *ARID1A* mutations, further suggesting that biallelic disruption is common. However, given the uncertainty of clonality estimates from single region data, the prevalence of biallelic truncal inactivation will need confirmation by MSeq in larger series. *ARID1A*-deficiency sensitizes cancer cells to small molecule inhibitors of the ATR DNA damage sensor^42^. Such a potential synthetic lethal interaction should be investigated in dMMR GOAs. Additional subclonal mutations in *ARID1A* and in other SWI/SNF-complex members evolved during cancer progression, indicating a role of SWI/SNF-complex modulation during carcinogenesis *and* cancer progression. MSeq and single sample TCGA data analysis also showed that chromosome 8 gains are among the earliest genetic events in ∼60% of these tumours. Further studies are necessary to investigate whether this is relevant for the tolerance of the MSI phenotype or a marker of aggressiveness as described for other cancer types^43,44^.

Taken together, the dMMR phenotype remained active throughout the evolution of primary tumours and in metastatic sites, generating extreme ITH. We furthermore revealed the generation of multiple subclonal driver mutations, including remarkable parallel evolution of multiple functionally similar subclonal drivers and a dN/dS ratio indicating positive selection in together 3 of 4 tumours. These results confirm a high evolvability of dMMR tumours. High heterogeneity and evolvability are thought to enable cancer aggressiveness and poor outcomes^45^, yet these data demonstrate a paradoxical association with good prognosis in dMMR tumours. dMMR tumours are unique models to advance insights into cancer evolution rules and into the potential and current limitations of evolutionary metrics for clinical outcome prediction.

## Methods

### Sample Collection and Preparation

Samples from treatment-naïve GOA resection specimens were routinely paraffin embedded and fresh frozen at the University Medical Center Hamburg-Eppendorf (Germany). The research use of specimens left over after the pathological diagnosis has been made is regulated through the ‘Hamburger Krankenhausgesetz’ (Hamburg Hospital Law) in Hamburg and consent and ethical approval are explicitly waived for samples that are fully anonymised. Thus, information about age, sex of the patients and outcome data is not available. The head of the Hamburg ethics committee confirmed in writing that no further ethics approval is required for this study.

Immunohistochemistry for MLH1, PMS2, MSH2 and MSH6 was performed on 20 cases and 4 with dMMR (each showing absence of MLH1 and PMS2 staining in cancer cells, see Fig. 1B) were identified by a pathologist. Seven tumour regions representing the spatial extent of each primary tumour were selected (surface area ∼8 × 5 mm and a depth of ∼10 mm) based on the H&E slide and spatial location within the tumour by a pathologist. Two cases each included two lymph node metastases (Station 1 – 2, right and left paracardial nodes) which were sufficiently large for analysis. Where necessary, samples were macrodissected to minimize stromal contamination. DNA was extracted using the Qiagen AllPrep kit following the manufacturer’s instructions. Nucleic acid yields were determined by Qubit (Invitrogen), and the quality and integrity of DNA was examined by agarose gel electrophoresis. DNA from tumour adjacent non-malignant tissue was used as a source of normal (‘germline’) DNA. For this either oesophageal or gastric wall tissue, embedded as “normal mucosa” was chosen and tumour contamination excluded by a pathologist based on H&E slides taken from levels before and after slides for DNA extraction.

### Multiplex Immunohistochemistry

The Opal 7 multiplexed assay (PerkinElmer) was used to perform combined CD8 (cytotoxic T-cells), pan-Cytokeratin (tumour cells) and DAPI (counter-) stains for each region. In Tumour 2 two regions had not enough tissue left after DNA extraction. Slides were scanned using the Vectra 3.0 pathology imaging system (PerkinElmer) as described before^46^.

After low magnification scanning, intratumour regions of interest were scanned at high resolution (20x). Spectral unmixing, tissue and cell segmentation and phenotyping of CD8 and Cytokeratin positive cells were performed with InForm image analysis software under pathologist supervision. Five representative regions of interests were chosen and cytotoxic T-cells and tumour cells in cancer tissue segmented areas were quantified. From the sum of the five regions we calculated the ratio of cytotoxic T-cells/tumour cells for each region of Tumours 1 -4.

### Whole exome sequencing

Tumour and matched germline DNA were sequenced by the NGS-Sequencing facility of the Tumour Profiling Unit at the Institute of Cancer Research. Exome sequencing libraries were prepared from 1 µg DNA using the Agilent SureSelectXT Human All Exon v6 kit according to the manufacturer’s protocol. Paired-end sequencing was performed on the Illumina HiSeq 2500 or NovaSeq 6000 with a minimum target depth of 100X in the adjacent normal samples and a minimum target depth of 200X in tumour regions.

BWA-MEM ^47^ (v0.7.12) was used to align the paired-end reads to the hg19 human reference genome to generate BAM format files. Picard Tools (http://picard.sourceforge.net) (v2.1.0) MarkDuplicates was run with duplicates removed. BAM files were coordinate sorted and indexed with SAMtools ^48^ (v0.1.19). BAM files were quality controlled using GATK ^49^ (v3.5-0) DepthOfCoverage, Picard CollectAlignmentSummaryMetrics (v2.1.0) and fastqc (https://www.bioinformatics.babraham.ac.uk/projects/fastqc/) (v0.11.4).

### Somatic mutation analysis

Single nucleotide variant (SNV) calls were generated with MuTect^50^ (v1.1.7) and VarScan2 ^51^ (v2.4.1) and mutation calls from both callers were combined. MuTect was run with default settings and post-filtered for a minimum variant frequency of 2%. SNVs generated by MuTect and flagged with ‘PASS’, ‘alt_allele_in_normal’ or ‘possible_contamination’ were retained. SAMtools (v1.3) mpileup was run with minimum mapping quality 1 and minimum base quality 20. The pileup file was inputted to VarScan2 somatic and run with a minimum variant frequency of 2%. The VarScan2 call loci were converted to BED format and bam-readcount (https://github.com/genome/bam-readcount) (v0.7.4) run on these positions with minimum mapping quality 1. The bam-readcount output allowed the VarScan2 calls to be further filtered using the recommended fpfilter.pl accessory script^52^ run on default settings. Indel calls were generated using Platypus^53^ (v.0.8.1) callVariants run on default settings. Calls were filtered based on the following FILTER flags - ‘GOF, ‘badReads, ‘hp10,’ MQ’, ‘strandBias’,’ QualDepth’,’ REFCALL’. We then filtered for somatic indels with normal genotype to be homozygous, minimum depth >= 10 in the normal, minimum depth >=20 in the tumour and >= 5 variant reads in the tumour.

The bam-readcount tool was run on all SNV loci using minimum mapping quality 1 and minimum base quality 5 to generate call QC metrics (e.g. average variant base quality, average variant mapping quality). High confidence SNVs were identified by filtering minimum average variant mapping quality 55 and minimum average variant base quality 35 in called tumour regions based on the bam-readcount QC metrics. Bam-readcount was then run on the filtered loci using minimum mapping quality 10 and minimum base quality 20 to generate allele counts for the merged VarScan2 and MuTect call loci. All SNV and indel calls were required to have a depth of at least 70 across all tumour regions. SNVs at positions sequenced to less than 20x depth in the matched germline and those which showed a variant frequency in the germline >2% and a variant count >2 were also excluded. Retained mutation calls were then passed through a cross-’germline’ filter that flags SNV and indel calls which are present with a VAF of >=2% in one of fourteen normal samples from the same sample collection. A call is rejected if the variant is flagged as present in 20% or more of the normal samples to remove common alignment artefacts or those arising recurrently at genomic positions which are difficult to sequence. Finally, calls were retained if at least one tumour region had a variant frequency of at least 5% and the same call was reported in all other regions of the same tumour showing a variant frequency of at least 2.5%. Variant calls on chromosomes X and Y were not considered.

SNV and indel calls were annotated using annovar ^54^ (v20160201) and oncotator ^55^ (v1.8.0.0 and oncotator_v1_ds_Jan262015 database) with hg19 build versions. The oncotator ‘COSMIC_n_overlapping_mutations’ field was used to flag mutations as possible drivers if they occurred in oncogenes and tumour suppressor genes in the online COSMIC Cancer Gene Census (CGC)^56^ or in driver genes identified in MSI tumours in the TCGA STAD publication ^2^. Mutations were defined as likely driver genes if they led to 1) an amino acid alteration that had previously been described in the COSMIC database 2) a disrupting mutation, including frameshift-, splice site- or premature stop/nonsense-mutations in a tumour suppressor gene or 3) an amino acid alteration at a position that shows an alteration in the COSMIC CGC but is distinct from the change reported in COSMIC if it was considered a likely driver by the Cancer Genome Interpreter^57^.

### DNA copy number aberration analysis

CNVKit^58^ (v0.8.1) was run in non-batch mode for copy number evaluation. Basic target and antitarget files were generated based on the Agilent SureSelectXT Human All Exon v6 kit. Accessible regions suggested by CNVKit (provided in the source distribution as ‘access-5kb-mappable.hg19.bed’) with a masked HLA interval (chr6:28866528-33775446) form the accessible loci. A pooled normal sample was created from all sequenced germline samples in the series. The copynumber^59^ R^60^ library functions Winsorize (run with ‘return.outliers’=TRUE) and pcf (run with ‘gamma’=200) were used to identify outliers and regions of highly uneven coverage (defined as an absolute log ratio value greater than 0.5) to exclude from the analysis.

We identified high confidence SNP locations using bcftools call^48^ (v1.3) with snp137 reference and SnpEff SnpSift^61^ (v4.2) to filter heterozygous loci with minimum depth 50. VarScan2 was used to call the tumour sample BAMs at these locations to generate B-Allele Frequency (BAF) data as input for CNVKit. CNVKit was run with matched germline samples along with the adjusted access and antitarget files. For the segmentation step we ran the copynumber function pcf with gamma=70. Breakpoints were then fed into Sequenza^62^ (v2.1.2) to calculate estimates of purity/ploidy and these values were used to recenter and scale the LogR profiles in CNVKit. BAF and LogR profiles were also manually reviewed by two researchers to determine their likely integer copy number states. Adjustments were made in cases where both manual reviews identified a consensus solution that differed from the bioinformatically generated integer copy number profile.

### Cancer cell content, ploidy estimation and wGII

Cancer cell content was estimated using the scaling factor of the copy number consensus solution. Ploidy was estimated as follows,

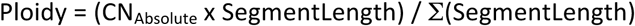

with CN_Absolute_ representing the unrounded copy number estimate and SegmentLength the genomic length between segment break points.

The weighted genome integrity index (wGII)^32^ is used to define chromosomal instability (CIN). We calculated the percentage of integer copy number segments on each chromosome different from the ploidy estimate rounded to the nearest integer state. The percentages are then averaged over the 22 autosomal chromsomes to give the wGII score.

### Subclonality analysis and phylogenetic tree reconstruction

Allele specific copy number estimates for SNV and indels were calculated as described in ^63^ and the cancer cell fraction (CCF) was estimated using the approach in ^64^. LICHeE^65^ was applied to infer phylogenetic trees from the estimated CCF values. The *build* algorithm was run with CCF/2 as input, -maxVAFAbsent 0, -minVAFPresent 0.0001 and ‘-s 10’. In each case we report the top ranked tree solution. A single valid tree was identified for Tumour 1 (error score: 0.02), Tumour 2 (error score: 0.13) and Tumour 4 (error score: 0.06). LICHeE identified 6 valid trees for Tumour 3 (error scores: 0.088, 0.095, 0.96, 0.106, 0.112, 0.113). These solutions differed only in the positioning of the branch immediately preceding H2 (which could be positioned at H1 or H3) and of that preceding G2 (which could be positioned at G1) in Fig. 3. The tree with the lowest error score was chosen for the analysis but selecting any of the alternative solutions would not change any of the the conclusions presented in this study. Otherwise, only a low percentage of mutations (2-7% per case) could not be assigned to a subclone in the phylogenetic tree

Trees were re-drawn and branch lengths scaled to the number of mutations in each subclonal mutation cluster and likely driver mutations were mapped onto the trunk or the appropriate branch. Private mutations identified by LICHeE were split into clonal and subclonal mutations using a CCF threshold of 0.7 unless the algorithm had already identified and split clonal and subclonal clusters. A short branch was added to Tumour 2 following a manual review of the tree solution to represent an 8 mutation cluster that was too small for the algorithm to detect but contained a *B2M* frameshift mutation which was identified as a likely driver.

### Mutational signatures

All SNV calls were loaded into R using VRanges (v1.28.3 ^66^ VariantAnnotation, given trinucleotide motifs using SomaticSignatures (v2.18.0)^67^ mutationContext and tabulated using motifMatrix with ‘normalize’=TRUE. The somatic motifs were then compared with the thirty mutational signatures established in COSMIC^68^ V2 using deconstructSigs(v1.8.0)^69^ whichSignatures and selecting ‘tri.counts.methods’=“exome”, ‘signature.cutoff’=0, ‘signatures.ref’=‘signatures.cosmic’ as run parameters. Mutation signatures representing at least 5% of mutations in one of the analysed mutation groups were reported.

#### Ratio of non-synonymous to synonymous mutations (dN/dS)

We ran dNdScv^38^ to generate dN/dS estimates which use trinucleotide context dependent substitution matrices to adjust for common mutation biases. We ran dNdScv with the following optional parameters: ‘outp = 1’, ‘max_muts_per_gene_per_sample = inf’ and ‘max_coding_muts_per_sample = inf’. This was done separately for mutations shown as truncal (blue), shared (yellow) and private (red and purple) on the phylogenetic trees in Fig. 3.

### HLA mutations and LOH calling

Mutations in HLA genes were predicted using the program POLYSOLVER^70^. In particular, we first predicted patients’ HLA types from germline samples using the shell_call_hla_type script of the POLYSOVER suite, with the following parameters: race=Unknown, includeFreq=1 and insertCalc=0. Then, we used these HLA predictions as input to the shell_call_hla_mutations_from_type script for predicting HLA mutations in tumour samples. Finally, the shell_annotate_hla_mutations script was used to annotate the mutations identified in the previous step.

Loss of heterozygosity events in HLA genes were predicted using the program LOHHLA^37^. LOHHLA requires as input normal HLA types, for which we used POLYSOLVER predictions, along with ploidy and cancer cell fraction estimates, which were available from the calculations described above. All other parameters were set to default values.

Neopeptides associated to somatic mutations were generated as decribed in ^71^. Note that we had to discard approximately 1.2% of somatic mutations because of inconsistencies between the variant annotation (this can be for either somatic variants or germline variants occurring on the same transcripts as the somatic ones) and the refseq_cds.txt file (GRCh37/hg19 Feb 2009) we used for generating the neopeptides. We used netMHCpan4.0 (28978689) to predict the neopeptides’ eluted ligand likelihood percentile rank scores. For each sample, we ran netMHCpan4.0 on all of the samples’ neopeptides against all samples’ HLA allotypes. As HLA-presented neopeptides we picked all *core* peptides (see ^71^) with a percentile rank < 0.5%.

### Comparison of non-silent mutation loads and clonal/subclonal drivers in TCGA MSI GOAs

64 GOAs from The Cancer Genome Atlas cohort^2^ are classified as MSI in the cBIO web portal^72^. We downloaded the BAM files of these cases from the NIH GDC Legacy Archive^73^. Adjustments to the analysis steps were necessary due to the properties of the TCGA sequencing data. A minimum variant frequency of 5% was applied throughout the mutation calling and the fpfilter.pl parameters ‘min-ref-avrl’ and ‘min-var-avrl’ filters were relaxed to 50. The minimum depth requirement in the tumour sample was relaxed to 20, while the minimum average base and mapping quality were set to 20 and 40 respectively. No adjustments were made to the default access and antitarget files of the CNVkit analysis due to large variations in the sequencing depths of the normal samples across the cohort. Otherwise, the somatic mutation, copy number and subclonality analysis steps were as described above. Mutational signatures were run as before and those detected with a mean contribution of 5% or more were further analysed.

## Data availability

Sequencing data have been deposited in the European Genome Phenome short read archive (Accession number: EGAS00001003434). Datasets are password protected and will be shared with researchers subject to signing a data sharing agreement.

## Acknowledgements

The study was supported by a Wellcome Trust Strategic Grant (105104/Z/14/Z) to the ICR Centre for Evolution and Cancer, by the National Institute for Health Research Biomedical Research Centre for Cancer at the ICR/RMH, by a Clinician Scientist Fellowship from Cancer Research UK and by grants from Cancer Genetics UK and the Constance Travis Trust.

## Author contributions

K.v.L. processed the tissue, designed and conducted experiment, analysed data and wrote the paper

A.W. performed bioinformatics analyses, analysed data and wrote the paper

M.P and S.L. performed bioinformatics analyses

B.G. processed the tissue

L.B., M.S., G.S. and B.C. analysed data

K.F., N.M. conducted exome sequencing

R.S., A.M., G.S. provided the tissue and performed dMMR analysis

M.G. designed the study, supervised the experiments and data analysis and wrote the paper

All authors read and approved the manuscript

## Declaration of Interests

The authors declare no competing interests.

## References

1 Ferlay, J. et al. Cancer incidence and mortality worldwide: sources, methods and major patterns in GLOBOCAN 2012. International journal of cancer. Journal international du cancer 136, E359–386, doi:10.1002/ijc.29210 (2015).

2 TCGA. Comprehensive molecular characterization of gastric adenocarcinoma. Nature, doi:10.1038/nature13480 (2014).

3 Polom, K. et al. Meta-analysis of microsatellite instability in relation to clinicopathological characteristics and overall survival in gastric cancer. The British journal of surgery 105, 159–167, doi:10.1002/bjs.10663 (2018).

4 Smyth, E. C. et al. Prognostic and predictive effect of microsatellite instability (MSI) in MAGIC. J Clin Oncol Suppl 3 (2015).

5 Cancer Genome Atlas Research, N. et al. Integrated genomic characterization of oesophageal carcinoma. Nature 541, 169–175, doi:10.1038/nature20805 (2017).

6 Frankell, A. M. et al. The landscape of selection in 551 esophageal adenocarcinomas defines genomic biomarkers for the clinic. Nature genetics 51, 506–516, doi:10.1038/s41588-018-0331-5 (2019).

7 Le, D. T. et al. Mismatch repair deficiency predicts response of solid tumors to PD-1 blockade. Science 357, 409–413, doi:10.1126/science.aan6733 (2017).

8 Kim, S. T. et al. Comprehensive molecular characterization of clinical responses to PD-1 inhibition in metastatic gastric cancer. Nature medicine 24, 1449–1458, doi:10.1038/s41591-018-0101-z (2018).

9 Gerlinger, M. et al. Cancer: Evolution Within a Lifetime. Annual review of genetics, doi:10.1146/annurev-genet-120213-092314 (2014).

10 Lipinski, K. A. et al. Cancer Evolution and the Limits of Predictability in Precision Cancer Medicine. Trends Cancer 2, 49–63 (2016).

11 Raynaud, F., Mina, M., Tavernari, D. & Ciriello, G. Pan-cancer inference of intra-tumor heterogeneity reveals associations with different forms of genomic instability. PLoS genetics 14, e1007669, doi:10.1371/journal.pgen.1007669 (2018).

12 Gerlinger, M. et al. Genomic architecture and evolution of clear cell renal cell carcinomas defined by multiregion sequencing. Nature genetics 46, 225–233, doi:10.1038/ng.2891 (2014).

13 Yap, T. A., Gerlinger, M., Futreal, P. A., Pusztai, L. & Swanton, C. Intratumor heterogeneity: seeing the wood for the trees. Science translational medicine 4, 127ps110, doi:10.1126/scitranslmed.3003854 (2012).

14 Albacker, L. A. et al. Loss of function JAK1 mutations occur at high frequency in cancers with microsatellite instability and are suggestive of immune evasion. PloS one 12, e0176181, doi:10.1371/journal.pone.0176181 (2017).

15 Shin, D. S. et al. Primary Resistance to PD-1 Blockade Mediated by JAK1/2 Mutations. Cancer discovery 7, 188–201, doi:10.1158/2159-8290.CD-16-1223 (2017).

16 de Bruin, E. C. et al. Spatial and temporal diversity in genomic instability processes defines lung cancer evolution. Science (New York, N.Y.) 346, 251–256, doi:10.1126/science.1253462 (2014).

17 Gerlinger, M. et al. Intratumor heterogeneity and branched evolution revealed by multiregion sequencing. The New England journal of medicine 366, 883–892, doi:10.1056/NEJMoa1113205 (2012).

18 McGranahan, N. & Swanton, C. Clonal Heterogeneity and Tumor Evolution: Past, Present, and the Future. Cell 168, 613–628, doi:10.1016/j.cell.2017.01.018 (2017).

19 Kawakami, H., Zaanan, A. & Sinicrope, F. A. Microsatellite instability testing and its role in the management of colorectal cancer. Curr Treat Options Oncol 16, 30, doi:10.1007/s11864-015-0348-2 (2015).

20 Harbst, K. et al. Multiregion Whole-Exome Sequencing Uncovers the Genetic Evolution and Mutational Heterogeneity of Early-Stage Metastatic Melanoma. Cancer research 76, 4765–4774, doi:10.1158/0008-5472.CAN-15-3476 (2016).

21 Alexandrov, L. B. et al. Signatures of mutational processes in human cancer. Nature 500, 415–421, doi:10.1038/nature12477 (2013).

22 McGranahan, N. et al. Clonal neoantigens elicit T cell immunoreactivity and sensitivity to immune checkpoint blockade. Science 351, 1463–1469, doi:10.1126/science.aaf1490 (2016).

23 Cristescu, R. et al. Pan-tumor genomic biomarkers for PD-1 checkpoint blockade-based immunotherapy. Science (New York, N.Y.) 362, doi:10.1126/science.aar3593 (2018).

24 McGranahan, N. et al. Clonal status of actionable driver events and the timing of mutational processes in cancer evolution. Sci Transl Med 7, 283ra254, doi:10.1126/scitranslmed.aaa1408 (2015).

25 Alexandrov, L. B. et al. Clock-like mutational processes in human somatic cells. Nature genetics 47, 1402–1407, doi:10.1038/ng.3441 (2015).

26 Blokzijl, F. et al. Tissue-specific mutation accumulation in human adult stem cells during life. Nature 538, 260–264, doi:10.1038/nature19768 (2016).

27 Yokoyama, A. et al. Age-related remodelling of oesophageal epithelia by mutated cancer drivers. Nature 565, 312–317, doi:10.1038/s41586-018-0811-x (2019).

28 Meier, B. et al. Mutational signatures of DNA mismatch repair deficiency in C. elegans and human cancers. Genome research 28, 666–675, doi:10.1101/gr.226845.117 (2018).

29 Dewhurst, S. M. et al. Tolerance of whole-genome doubling propagates chromosomal instability and accelerates cancer genome evolution. Cancer discovery 4, 175–185, doi:10.1158/2159-8290.CD-13-0285 (2014).

30 Birkbak, N. J. et al. Paradoxical relationship between chromosomal instability and survival outcome in cancer. Cancer research 71, 3447–3452, doi:10.1158/0008-5472.CAN-10-3667 (2011).

31 Pihan, G. & Doxsey, S. J. Mutations and aneuploidy: co-conspirators in cancer? Cancer cell 4, 89–94 (2003).

32 Burrell, R. A. et al. Replication stress links structural and numerical cancer chromosomal instability. Nature 494, 492–496, doi:10.1038/nature11935 (2013).

33 Giannakis, M. et al. RNF43 is frequently mutated in colorectal and endometrial cancers. Nat Genet, doi:10.1038/ng.3127 (2014).

34 Lewis, K. A. et al. Heterozygous ATR mutations in mismatch repair-deficient cancer cells have functional significance. Cancer research 65, 7091–7095, doi:10.1158/0008-5472.CAN-05-1019 (2005).

35 Gruber, S. B. et al. BLM heterozygosity and the risk of colorectal cancer. Science 297, 2013, doi:10.1126/science.1074399 (2002).

36 Wilson, B. G. & Roberts, C. W. SWI/SNF nucleosome remodellers and cancer. Nat Rev Cancer 11, 481–492, doi:10.1038/nrc3068 (2011).

37 McGranahan, N. et al. Allele-Specific HLA Loss and Immune Escape in Lung Cancer Evolution. Cell 171, 1259–1271 e1211, doi:10.1016/j.cell.2017.10.001 (2017).

38 Martincorena, I. et al. Universal Patterns of Selection in Cancer and Somatic Tissues. Cell 171, 1029–1041 e1021, doi:10.1016/j.cell.2017.09.042 (2017).

39 Zaretsky, J. M. et al. Mutations Associated with Acquired Resistance to PD-1 Blockade in Melanoma. The New England journal of medicine, doi:10.1056/NEJMoa1604958 (2016).

40 Sveen, A. et al. Multilevel genomics of colorectal cancers with microsatellite instability-clinical impact of JAK1 mutations and consensus molecular subtype 1. Genome Med 9, 46, doi:10.1186/s13073-017-0434-0 (2017).

41 Middha, S. et al. Majority of B2M-Mutant and -Deficient Colorectal Carcinomas Achieve Clinical Benefit From Immune Checkpoint Inhibitor Therapy and Are Microsatellite Instability-High. JCO Precis Oncol 3, doi:10.1200/PO.18.00321 (2019).

42 Williamson, C. T. et al. ATR inhibitors as a synthetic lethal therapy for tumours deficient in ARID1A. Nature communications 7, 13837, doi:10.1038/ncomms13837 (2016).

43 Steiner, T. et al. Gain in chromosome 8q correlates with early progression in hormonal treated prostate cancer. Eur Urol 41, 167–171 (2002).

44 Klatte, T. et al. Gain of chromosome 8q is associated with metastases and poor survival of patients with clear cell renal cell carcinoma. Cancer 118, 5777–5782, doi:10.1002/cncr.27607 (2012).

45 Maley, C. C. et al. Classifying the evolutionary and ecological features of neoplasms. Nat Rev Cancer 17, 605–619, doi:10.1038/nrc.2017.69 (2017).

46 Stack, E. C., Wang, C., Roman, K. A. & Hoyt, C. C. Multiplexed immunohistochemistry, imaging, and quantitation: a review, with an assessment of Tyramide signal amplification, multispectral imaging and multiplex analysis. Methods 70, 46–58, doi:10.1016/j.ymeth.2014.08.016 (2014).

47 Li, H. & Durbin, R. Fast and accurate short read alignment with Burrows-Wheeler transform. Bioinformatics (Oxford, England) 25, 1754–1760, doi:10.1093/bioinformatics/btp324 (2009).

48 Li, H. et al. The Sequence Alignment/Map format and SAMtools. Bioinformatics (Oxford, England) 25, 2078–2079, doi:10.1093/bioinformatics/btp352 (2009).

49 McKenna, A. et al. The Genome Analysis Toolkit: a MapReduce framework for analyzing next-generation DNA sequencing data. Genome research 20, 1297–1303, doi:10.1101/gr.107524.110 (2010).

50 Cibulskis, K. et al. Sensitive detection of somatic point mutations in impure and heterogeneous cancer samples. Nature biotechnology 31, 213–219, doi:10.1038/nbt.2514 (2013).

51 Koboldt, D. C. et al. VarScan 2: somatic mutation and copy number alteration discovery in cancer by exome sequencing. Genome research 22, 568–576, doi:10.1101/gr.129684.111 (2012).

52 Koboldt, D. C., Larson, D. E. & Wilson, R. K. Using VarScan 2 for Germline Variant Calling and Somatic Mutation Detection. Current protocols in bioinformatics 44, 15.14.11–17, doi:10.1002/0471250953.bi1504s44 (2013).

53 Rimmer, A. et al. Integrating mapping-, assembly- and haplotype-based approaches for calling variants in clinical sequencing applications. Nat Genet 46, 912–918, doi:10.1038/ng.3036 (2014).

54 Wang, K., Li, M. & Hakonarson, H. ANNOVAR: functional annotation of genetic variants from high-throughput sequencing data. Nucleic Acids Research 38, e164–e164, doi:10.1093/nar/gkq603 (2010).

55 Ramos, A. H. et al. Oncotator: cancer variant annotation tool. Human mutation 36, E2423–2429, doi:10.1002/humu.22771 (2015).

56 Sondka, Z. et al. The COSMIC Cancer Gene Census: describing genetic dysfunction across all human cancers. Nat Rev Cancer 18, 696–705, doi:10.1038/s41568-018-0060-1 (2018).

57 Tamborero, D. et al. Cancer Genome Interpreter annotates the biological and clinical relevance of tumor alterations. Genome Med 10, 25, doi:10.1186/s13073-018-0531-8 (2018).

58 Talevich, E., Shain, A. H., Botton, T. & Bastian, B. C. CNVkit: Genome-Wide Copy Number Detection and Visualization from Targeted DNA Sequencing. PLOS Computational Biology 12, e1004873, doi:10.1371/journal.pcbi.1004873 (2016).

59 Nilsen, G. et al. Copynumber: Efficient algorithms for single- and multi-track copy number segmentation. BMC Genomics 13, 591–591, doi:10.1186/1471-2164-13-591 (2012).

60 R: A Language and Environment for Statistical Computing (R Foundation for Statistical Computing, Vienna, Austria, 2018).

61 Cingolani, P. et al. A program for annotating and predicting the effects of single nucleotide polymorphisms, SnpEff: SNPs in the genome of Drosophila melanogaster strain w1118; iso-2; iso-3. Fly 6, 80–92, doi:10.4161/fly.19695 (2012).

62 Favero, F. et al. Sequenza: allele-specific copy number and mutation profiles from tumor sequencing data. Annals of oncology : official journal of the European Society for Medical Oncology 26, 64–70, doi:10.1093/annonc/mdu479 (2015).

63 Stephens, P. J. et al. The landscape of cancer genes and mutational processes in breast cancer. Nature 486, 400–404, doi:10.1038/nature11017 (2012).

64 Letouze, E. et al. Mutational signatures reveal the dynamic interplay of risk factors and cellular processes during liver tumorigenesis. Nature communications 8, 1315, doi:10.1038/s41467-017-01358-x (2017).

65 Popic, V. et al. Fast and scalable inference of multi-sample cancer lineages. Genome biology 16, 91, doi:10.1186/s13059-015-0647-8 (2015).

66 Obenchain, V. et al. VariantAnnotation: a Bioconductor package for exploration and annotation of genetic variants. Bioinformatics (Oxford, England) 30, 2076–2078, doi:10.1093/bioinformatics/btu168 (2014).

67 Gehring, J. S., Fischer, B., Lawrence, M. & Huber, W. SomaticSignatures: inferring mutational signatures from single-nucleotide variants. Bioinformatics (Oxford, England) 31, 3673–3675, doi:10.1093/bioinformatics/btv408 (2015).

68 Forbes, S. A. et al. COSMIC: somatic cancer genetics at high-resolution. Nucleic acids research 45, D777–D783, doi:10.1093/nar/gkw1121 (2017).

69 Rosenthal, R., McGranahan, N., Herrero, J., Taylor, B. S. & Swanton, C. DeconstructSigs: delineating mutational processes in single tumors distinguishes DNA repair deficiencies and patterns of carcinoma evolution. Genome biology 17, 31, doi:10.1186/s13059-016-0893-4 (2016).

70 Shukla, S. A. et al. Comprehensive analysis of cancer-associated somatic mutations in class I HLA genes. Nature biotechnology 33, 1152–1158, doi:10.1038/nbt.3344 (2015).

71 Woolston, A. et al. Genomic and Transcriptomic Determinants of Therapy Resistance and Immune Landscape Evolution during Anti-EGFR Treatment in Colorectal Cancer. Cancer cell 36, 35–50 e39, doi:10.1016/j.ccell.2019.05.013 (2019).

72 Gao, J. et al. Integrative analysis of complex cancer genomics and clinical profiles using the cBioPortal. Science signaling 6, pl1, doi:10.1126/scisignal.2004088 (2013).

73 Grossman, R. L. et al. Toward a Shared Vision for Cancer Genomic Data. The New England journal of medicine 375, 1109–1112, doi:10.1056/NEJMp1607591 (2016).

