## Supplementary Information for "Extreme intratumour heterogeneity and driver evolution in mismatch repair deficient gastro-oesophageal cancer"

#### Supplementary Material

#### Table of Contents

| | |
|---|---|
| Supplementary Figure 1. Clonal and subclonal neoantigen burden. | 2 |
| Supplementary Figure 2. Plots of somatic mutation counts split by the 96 mutation types. | 3 |
| Supplementary Figure 3. Allele specific copy number profiles of individual tumour regions. | 6 |
| Supplementary Figure 4. An illustration of phylogenetic conflict in Tumour 1. | 10 |
| Supplementary Figure 5. Mapping of phylogenetic trees onto tumour maps. | 11 |
| Supplementary Figure 6. Possible neoantigen loss due to HLA mutations. | 12 |
| Supplementary Figure 7. Paired-end reads demonstrate biallelic inactivation of *B2M*. | 13 |
| Supplementary Figure 8. Trunk overestimation by single region analysis. | 14 |
| Supplementary Figure 9. Mutation copy number of chromosome 8 in the STAD TCGA series. | 15 |
| Supplementary Table 1. Mseq mutations calls and depths. | 23 |
| Supplementary Table 2. Cell purity and ploidy. | 23 |
| Supplementary Table 3. COSMIC signatures. | 23 |
| Supplementary Table 4. Putative driver mutations. | 23 |
| Supplementary Table 5. HLA mutation calls and LOH analysis. | 23 |
| Supplementary Table 6. dN/dS. | 23 |

**Supplementary Figure 1. Clonal and subclonal neoantigen burden.**

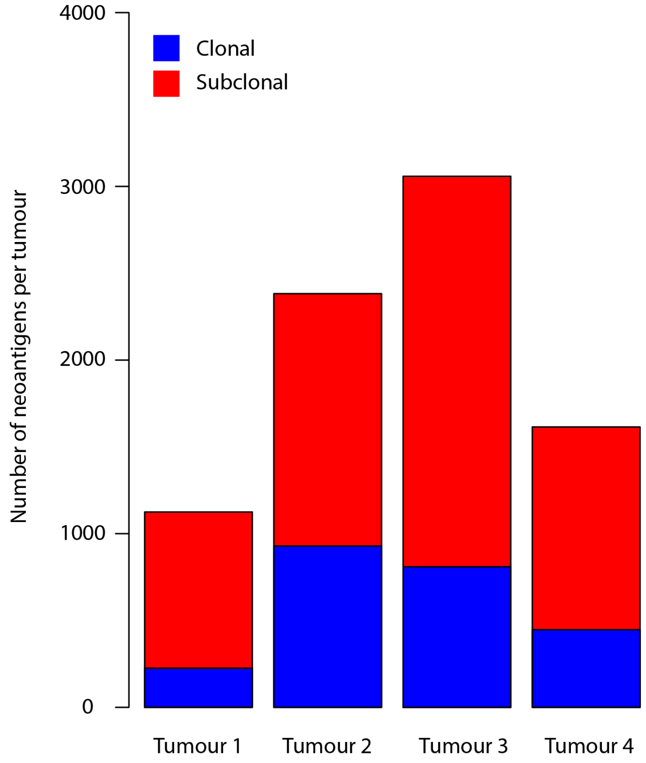

**Supplementary Figure 2. Plots of somatic mutation counts split by the 96 mutation types.**

**A**. Normalised mutation spectrums for Tumours 1-4. Plots are based on the following SNV burdens: Tumour 1: 1653, Tumour 2: 1766, Tumour 3: 4662 amd Tumour 4: 1738. **B**. Normalised mutation profiles split by private, shared and truncal mutations for each tumour. Plots are based on the following SNV burdens: Tumour 1: 835 private, 490 shared, 328 truncal, Tumour 2: 480 private, 495 shared, 791 truncal, Tumour 3: 2002 private, 1698 shared, 962 truncal, Tumour 4: 580 private, 660 shared, 498 truncal.

**A**

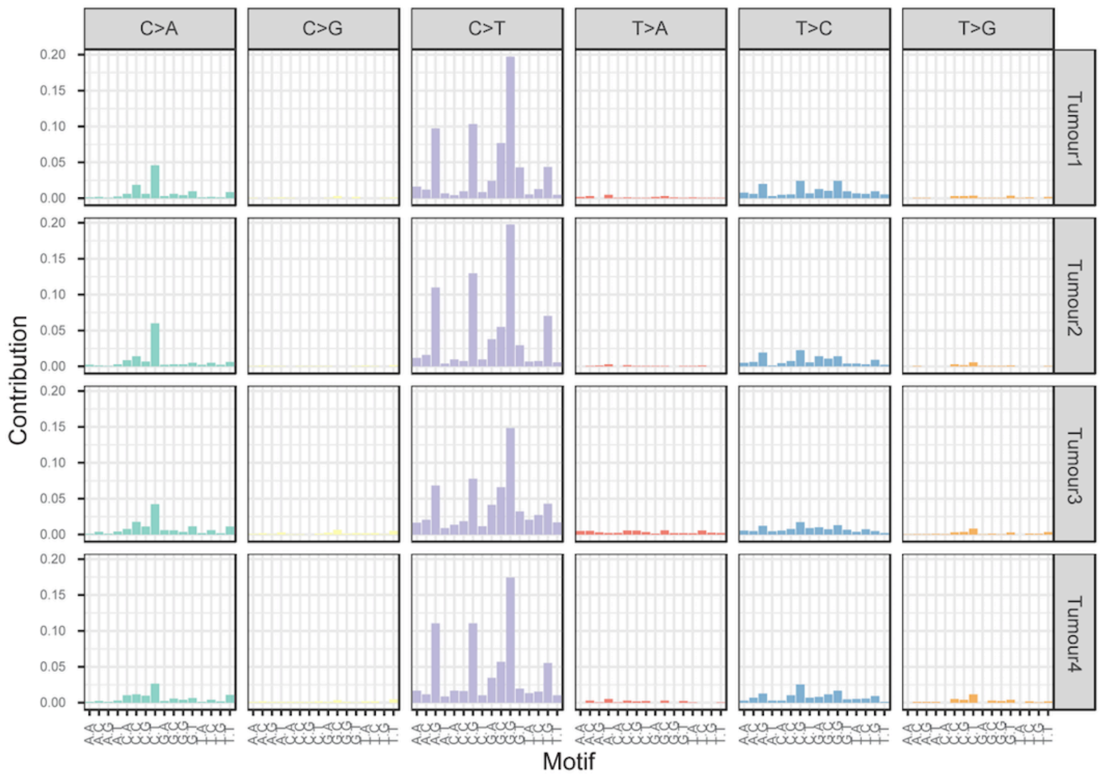

**B**

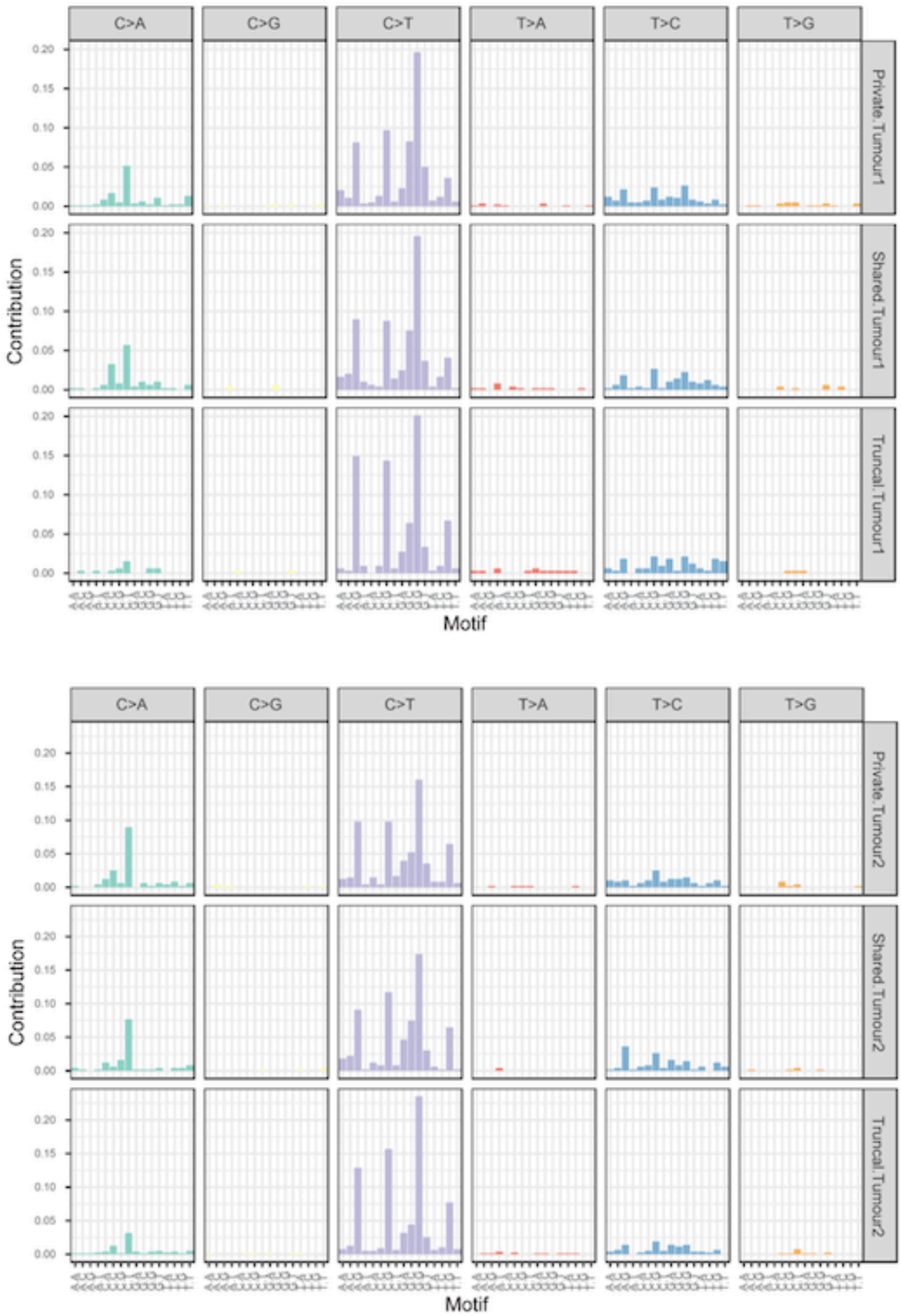
C>A
C>G
C>T
T>A
T>C
T>G
Private.Tumour1
Shared.Tumour1
Truncal.Tumour1
Contribution
0.20
0.15
0.10
0.05
0.00
Motif
C>A
C>G
C>T
T>A
T>C
T>G
Private.Tumour2
Shared.Tumour2
Truncal.Tumour2
Contribution
0.20
0.15
0.10
0.05
0.00
Motif

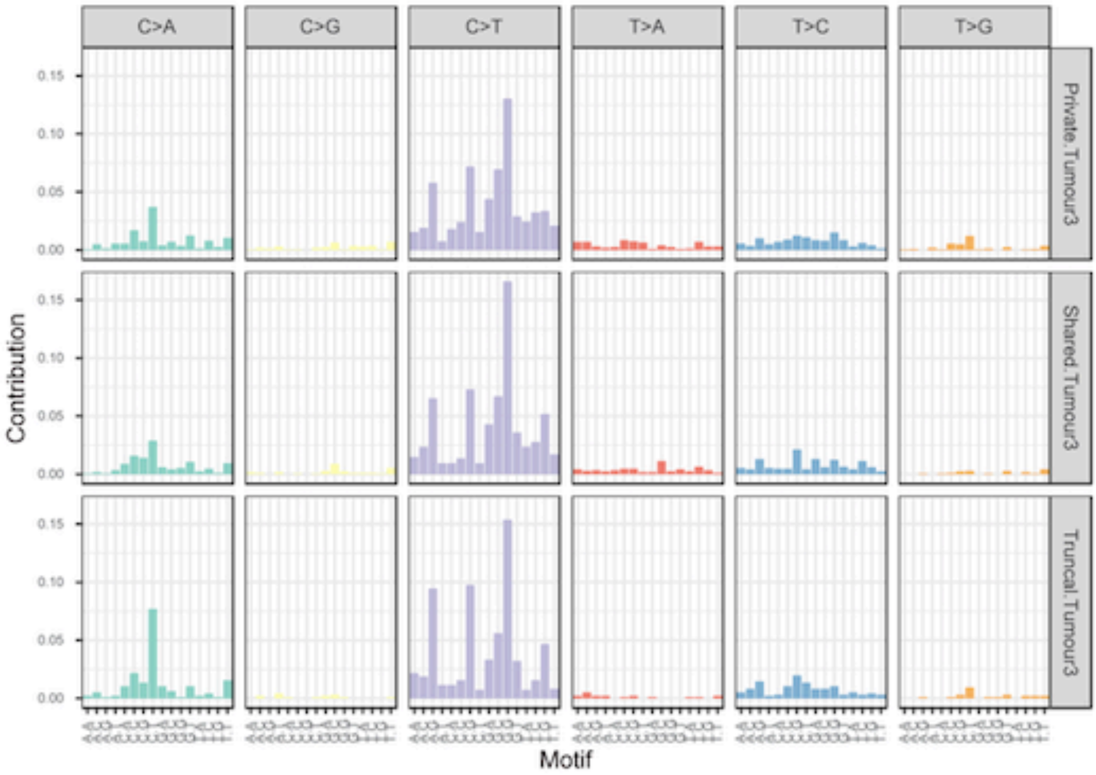
C>A
C>G
C>T
T>A
T>C
T>G
Private.Tumour3
Shared.Tumour3
Truncal.Tumour3
0.15
0.10
0.05
0.00
Contribution
Motif

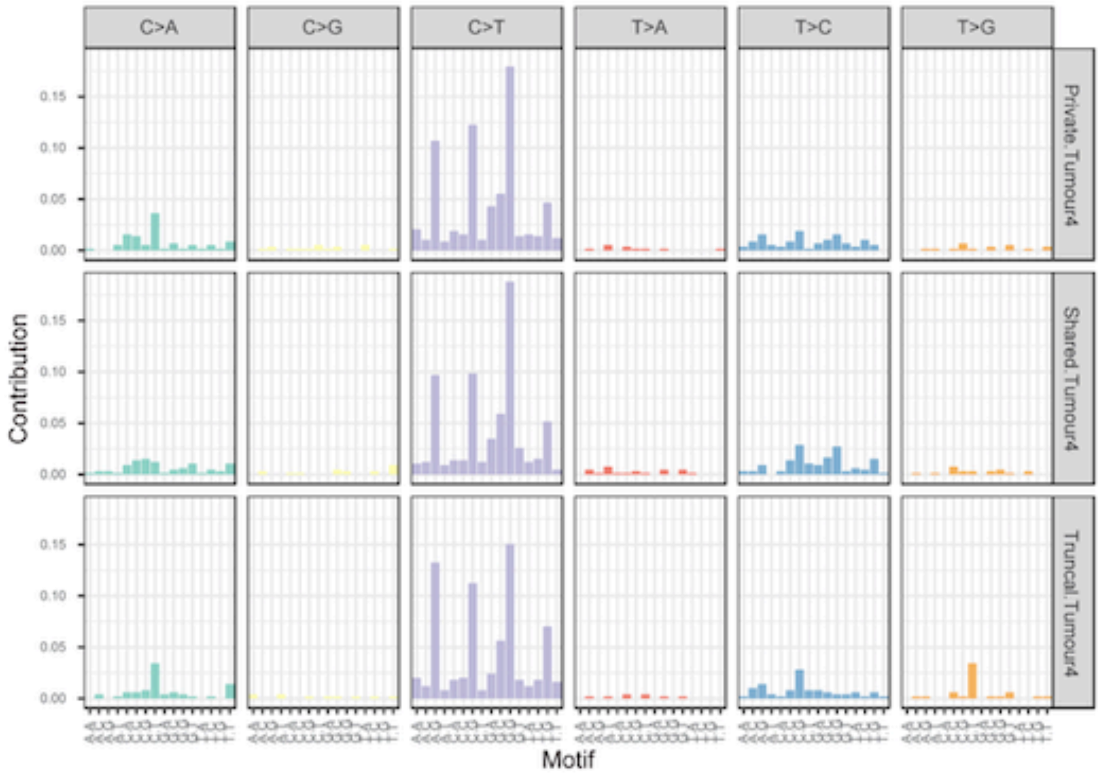
C>A
C>G
C>T
T>A
T>C
T>G
Private.Tumour4
Shared.Tumour4
Truncal.Tumour4
0.15
0.10
0.05
0.00
Contribution
Motif

### Supplementary Figure 3. Allele specific copy number profiles of individual tumour regions

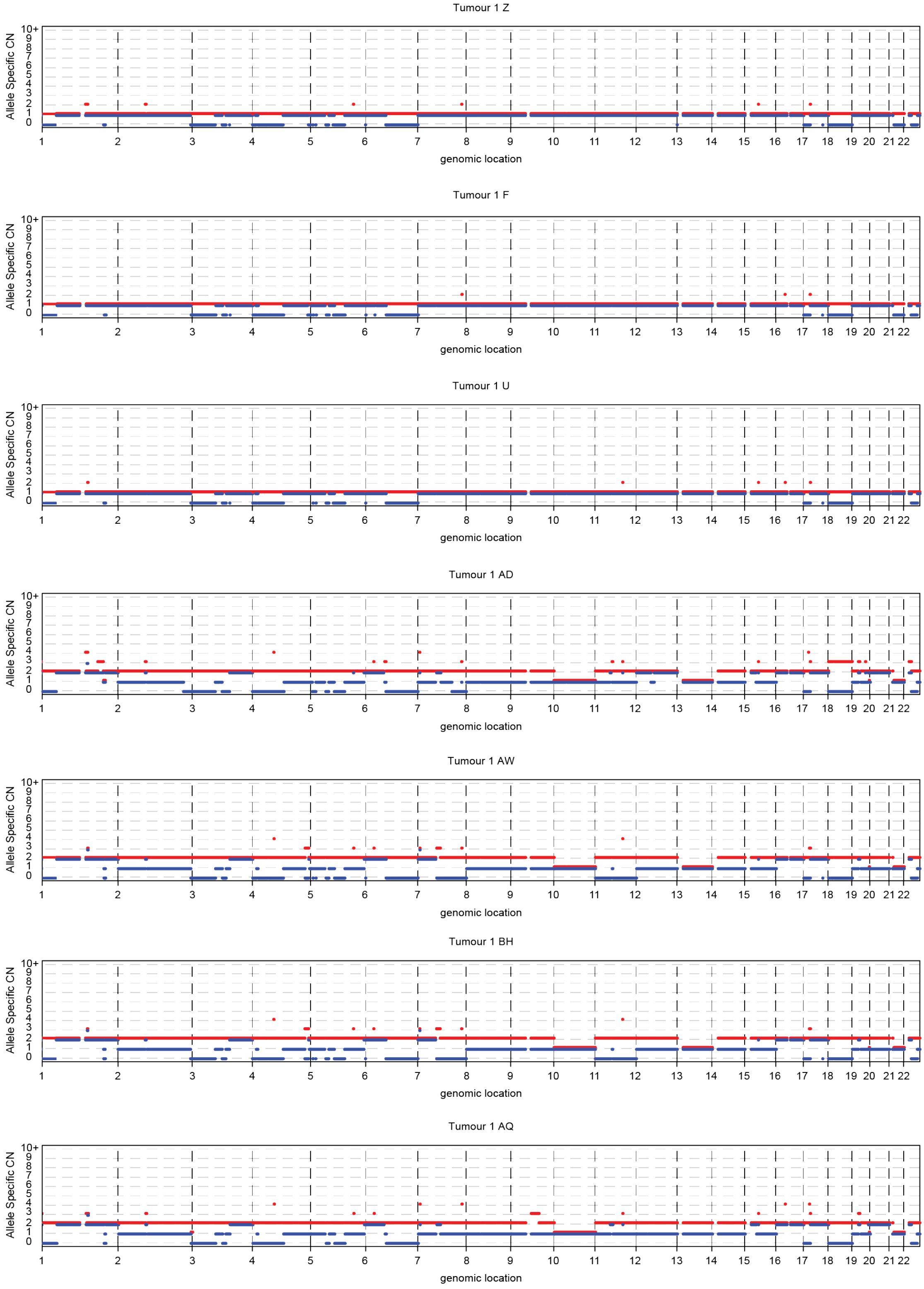

Tumour 2 E

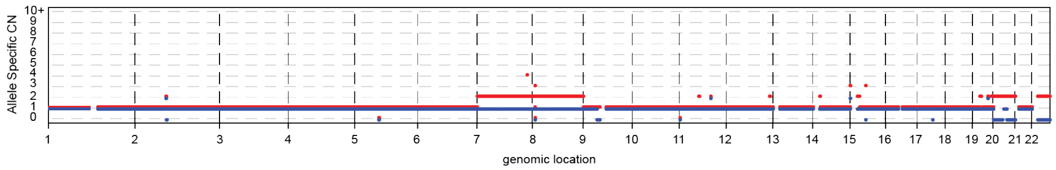

Tumour 2 T

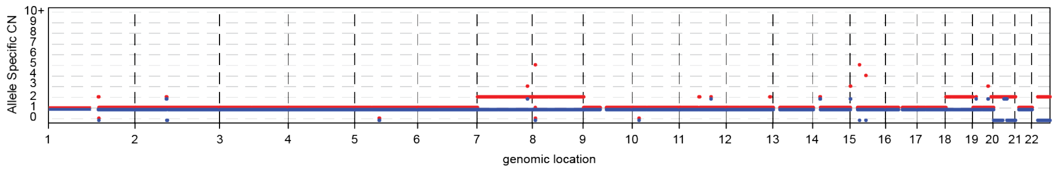

Tumour 2 P

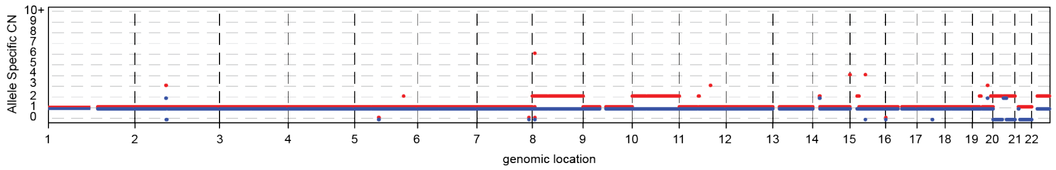

Tumour 2 Y

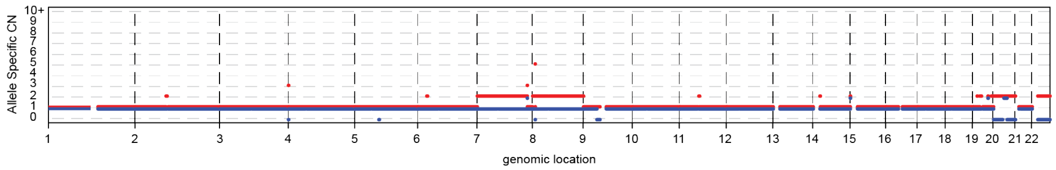

Tumour 2 S

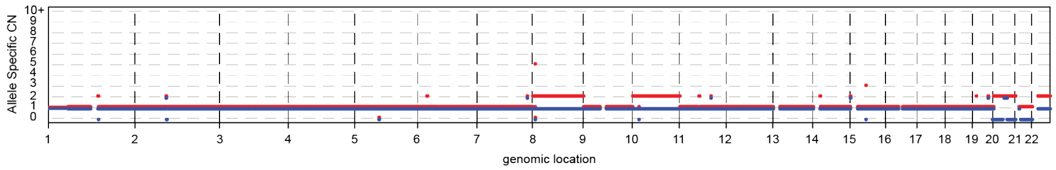

Tumour 2 N

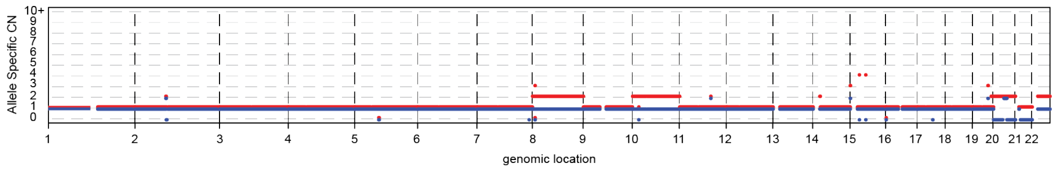

Tumour 2 AE

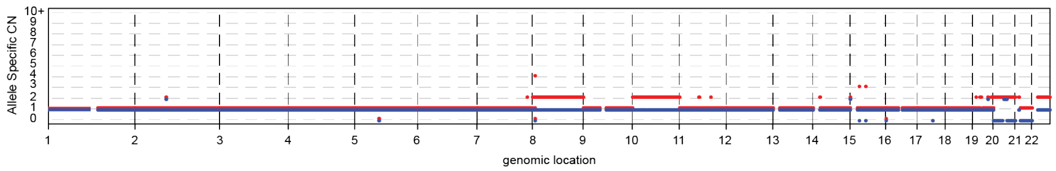

Tumour 2 AK

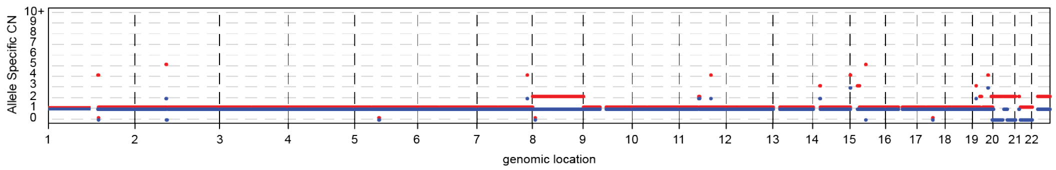

Tumour 2 AL

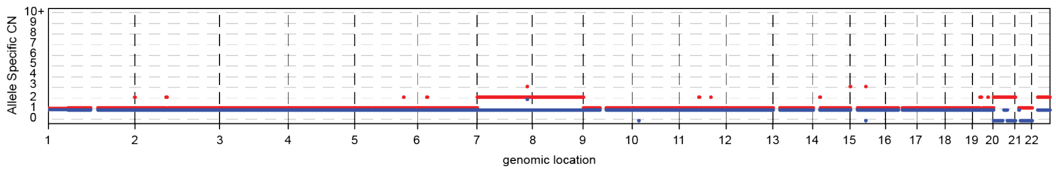

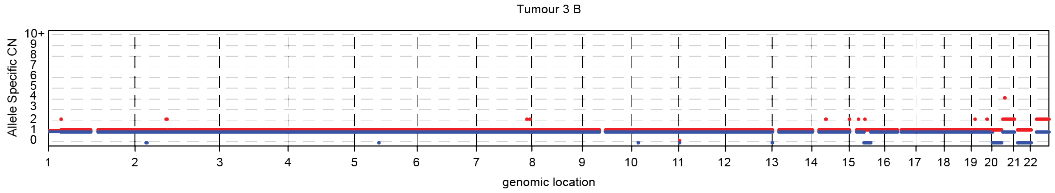
Tumour 3 B
Allele Specific CN
10+ 9 8 7 6 5 4 3 2 1 0
1 2 3 4 5 6 7 8 9 10 11 12 13 14 15 16 17 18 19 20 21 22
genomic location

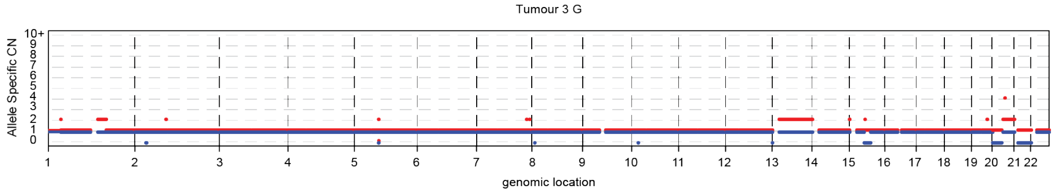
Tumour 3 G
Allele Specific CN
10+ 9 8 7 6 5 4 3 2 1 0
1 2 3 4 5 6 7 8 9 10 11 12 13 14 15 16 17 18 19 20 21 22
genomic location

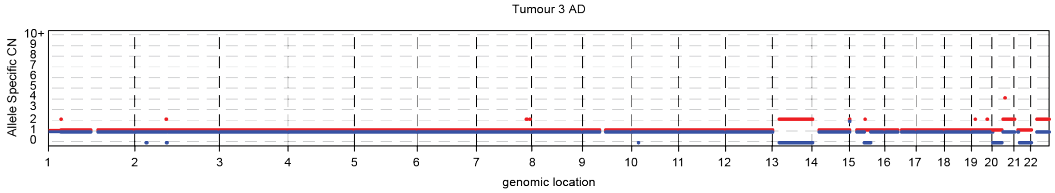
Tumour 3 AD
Allele Specific CN
10+ 9 8 7 6 5 4 3 2 1 0
1 2 3 4 5 6 7 8 9 10 11 12 13 14 15 16 17 18 19 20 21 22
genomic location

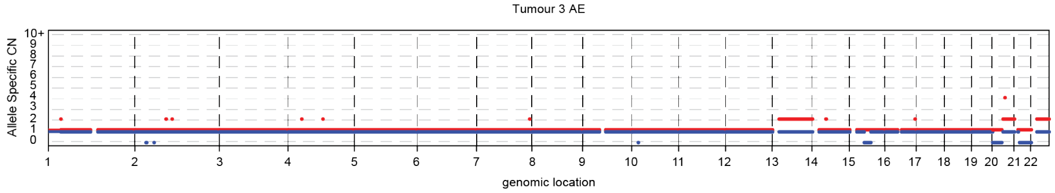
Tumour 3 AE
Allele Specific CN
10+ 9 8 7 6 5 4 3 2 1 0
1 2 3 4 5 6 7 8 9 10 11 12 13 14 15 16 17 18 19 20 21 22
genomic location

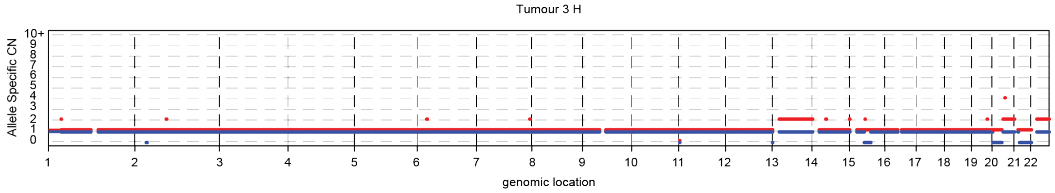
Tumour 3 H
Allele Specific CN
10+ 9 8 7 6 5 4 3 2 1 0
1 2 3 4 5 6 7 8 9 10 11 12 13 14 15 16 17 18 19 20 21 22
genomic location

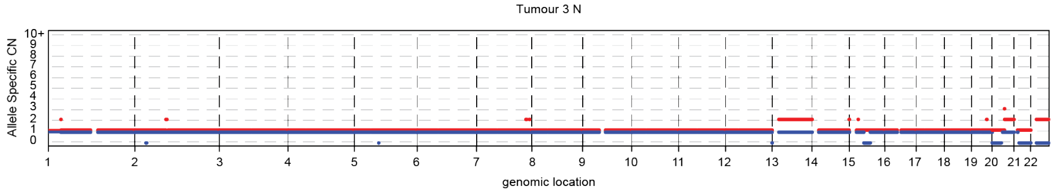
Tumour 3 N
Allele Specific CN
10+ 9 8 7 6 5 4 3 2 1 0
1 2 3 4 5 6 7 8 9 10 11 12 13 14 15 16 17 18 19 20 21 22
genomic location

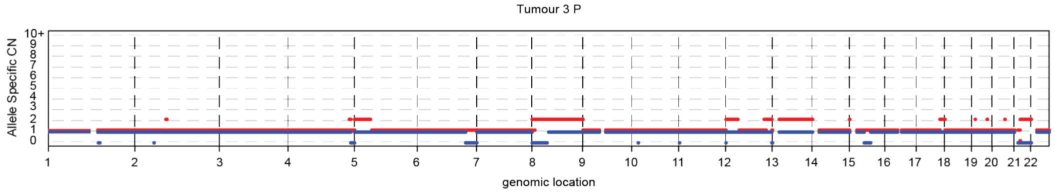
Tumour 3 P
Allele Specific CN
10+ 9 8 7 6 5 4 3 2 1 0
1 2 3 4 5 6 7 8 9 10 11 12 13 14 15 16 17 18 19 20 21 22
genomic location

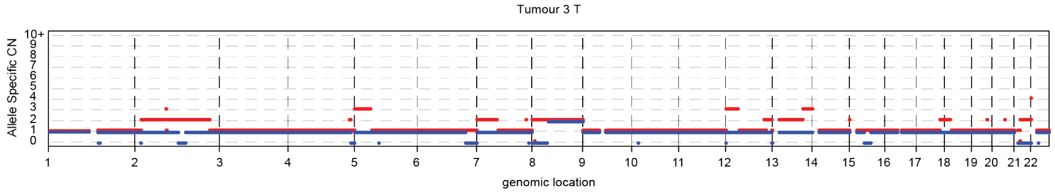
Tumour 3 T
Allele Specific CN
10+ 9 8 7 6 5 4 3 2 1 0
1 2 3 4 5 6 7 8 9 10 11 12 13 14 15 16 17 18 19 20 21 22
genomic location

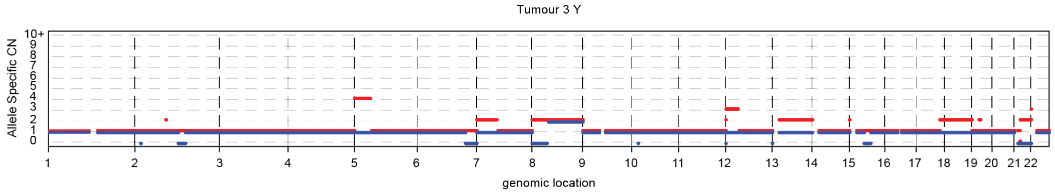
Tumour 3 Y
Allele Specific CN
10+ 9 8 7 6 5 4 3 2 1 0
1 2 3 4 5 6 7 8 9 10 11 12 13 14 15 16 17 18 19 20 21 22
genomic location

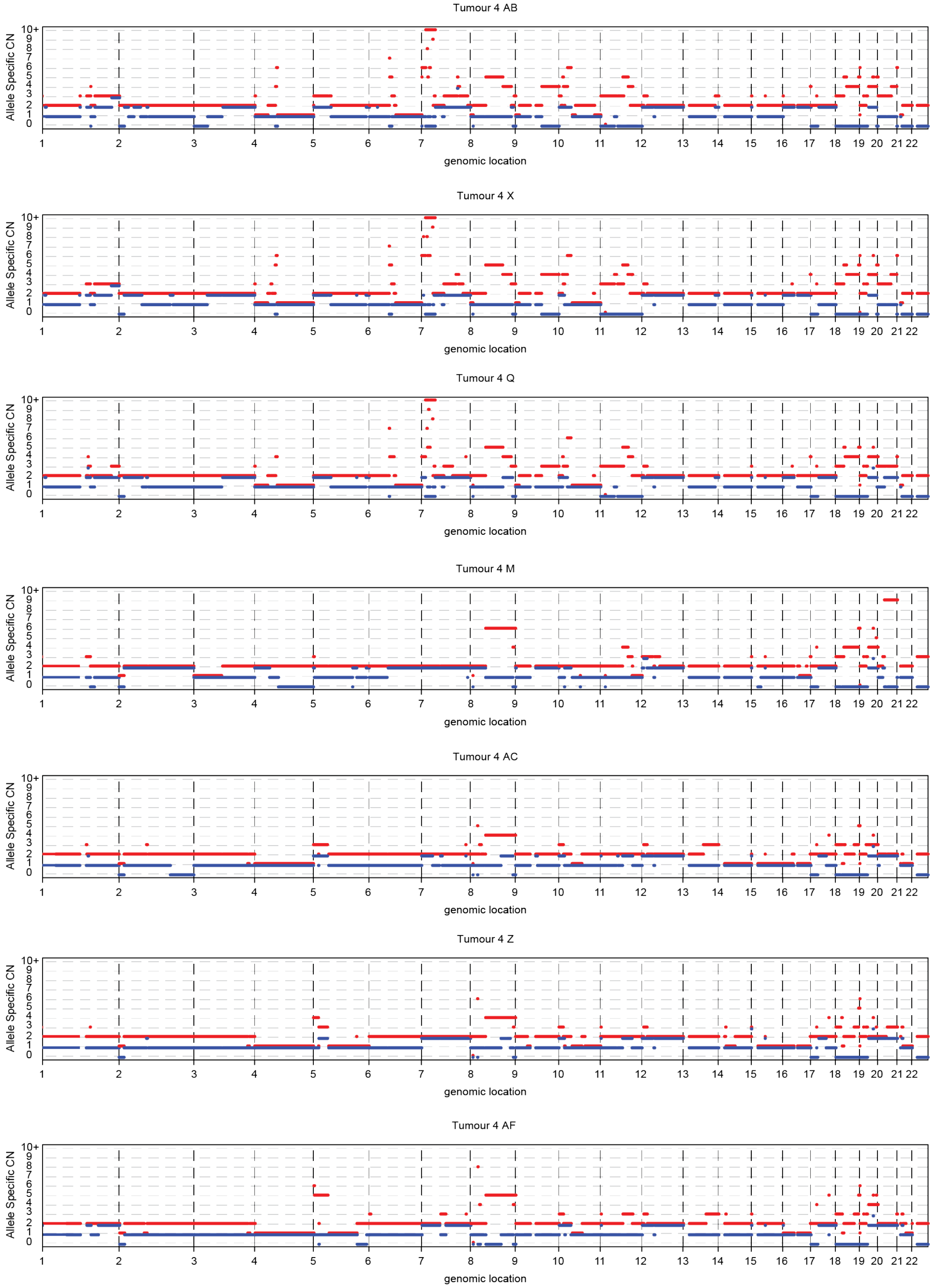
Tumour 4 AB
Tumour 4 X
Tumour 4 Q
Tumour 4 M
Tumour 4 AC
Tumour 4 Z
Tumour 4 AF
Allele Specific CN
genomic location
10+
1 2 3 4 5 6 7 8 9 10 11 12 13 14 15 16 17 18 19 20 21 22

**Supplementary Figure 4. An Illustration of Phylogenetic conflict in Tumour 1.**

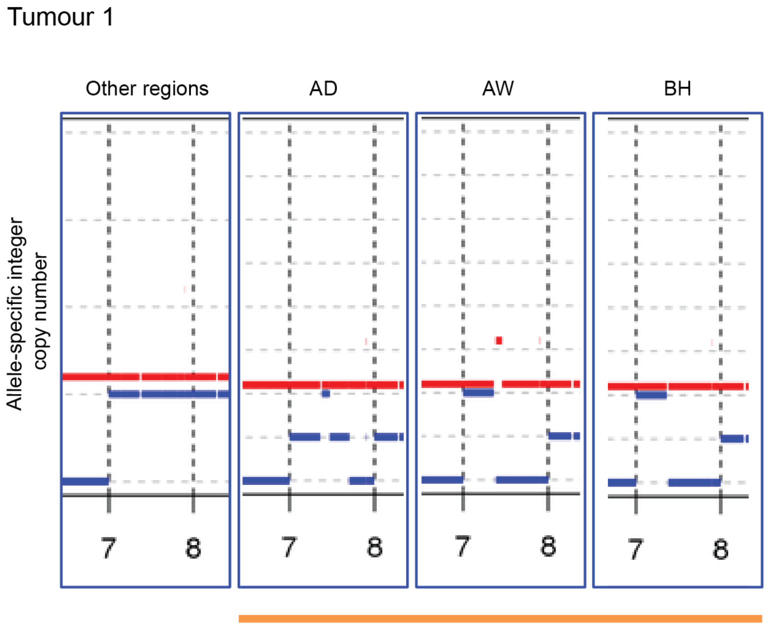

Chr7q LOH in AW and BH and shorter segment LOH in AD which explain one phylogenetic conflict in Tumour 1

#### Supplementary Figure 5. Mapping of phylogenetic trees onto tumour maps.

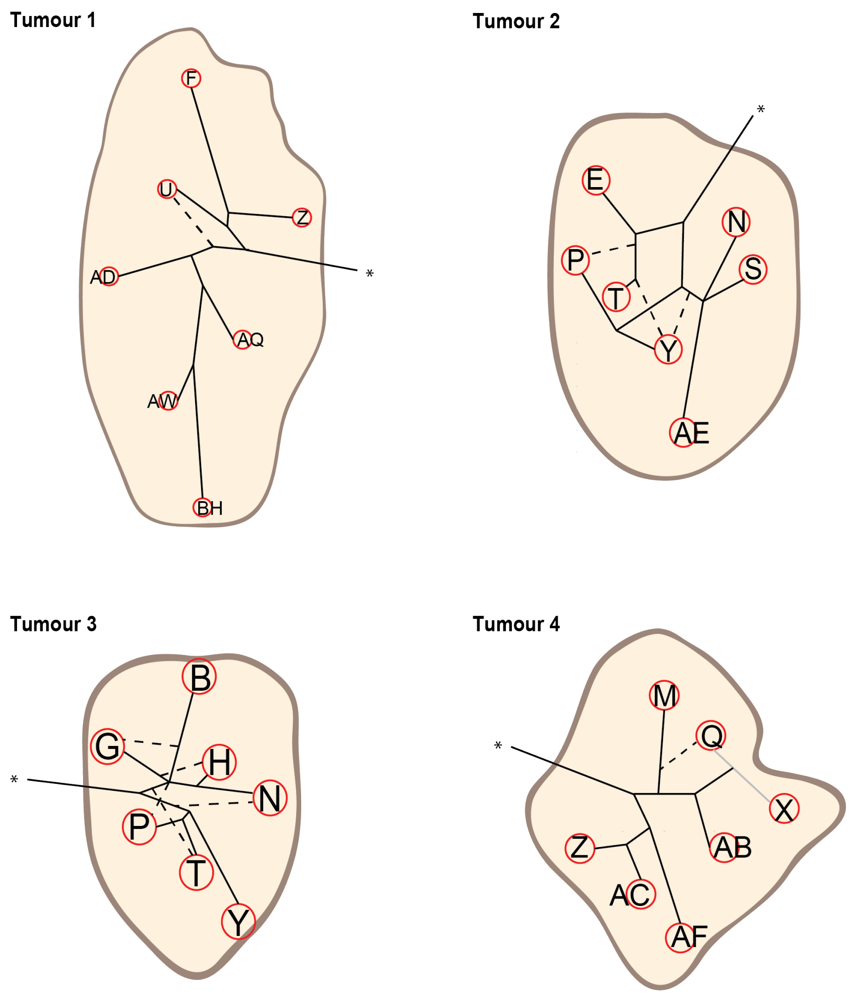

Phylogenetic tree trunk or branches indicating ancestral relationships of major clones
Phylogenetic tree branches indicating ancestral relationships of minority subclones
Connecting line indicating subclones which appeared identical in the phylogenetic deconvolution
Trunk and branch lengths are scale free

*Origin of the phylogenetic tree, mapped outside of the tumour area by default

#### Supplementary Figure 6. Possible neoantigen loss due to HLA mutations.

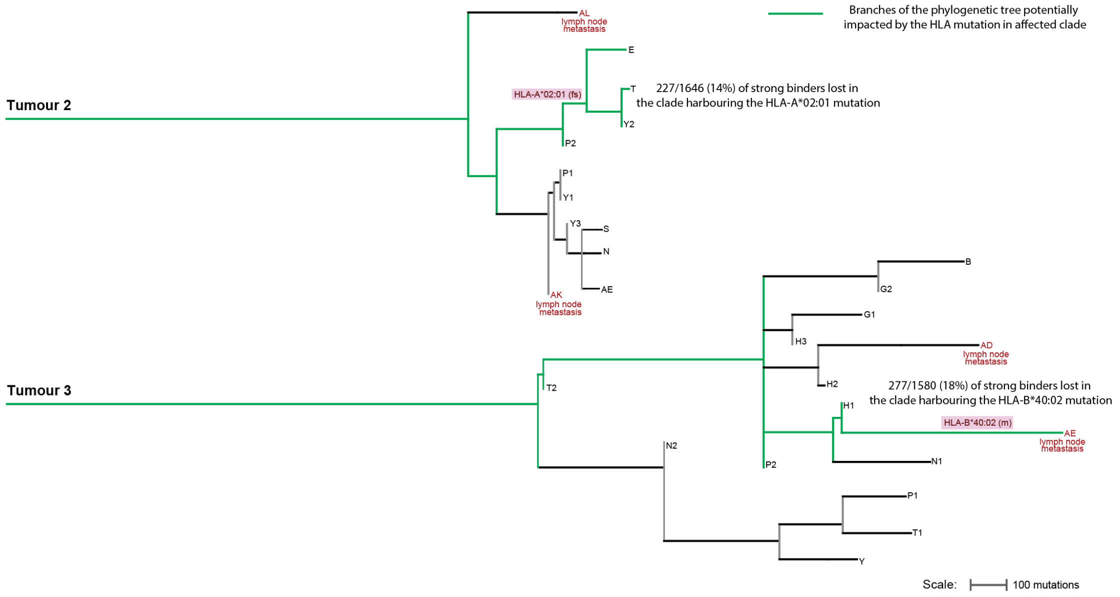

#### Supplementary Figure 7. Paired-end reads demonstrate biallelic inactivation of *B2M*.

Screenshots from the Interactive Genome Viewer of Tumour 2 AE displaying the *B2M* splice-site and frameshift mutations. Paired reads that cover both mutation loci are highlighted with red boxes. **A.** A total of 4 read pairs showed the *B2M* splice-site mutation and covered the location of the *B2M* frameshift mutation but none showed the frameshift mutation. **B.** A total of ten read pairs showed the *B2M* frameshift mutation and covered the location of the *B2M* splice site mutations but none showed the *B2M* splice-site mutation.

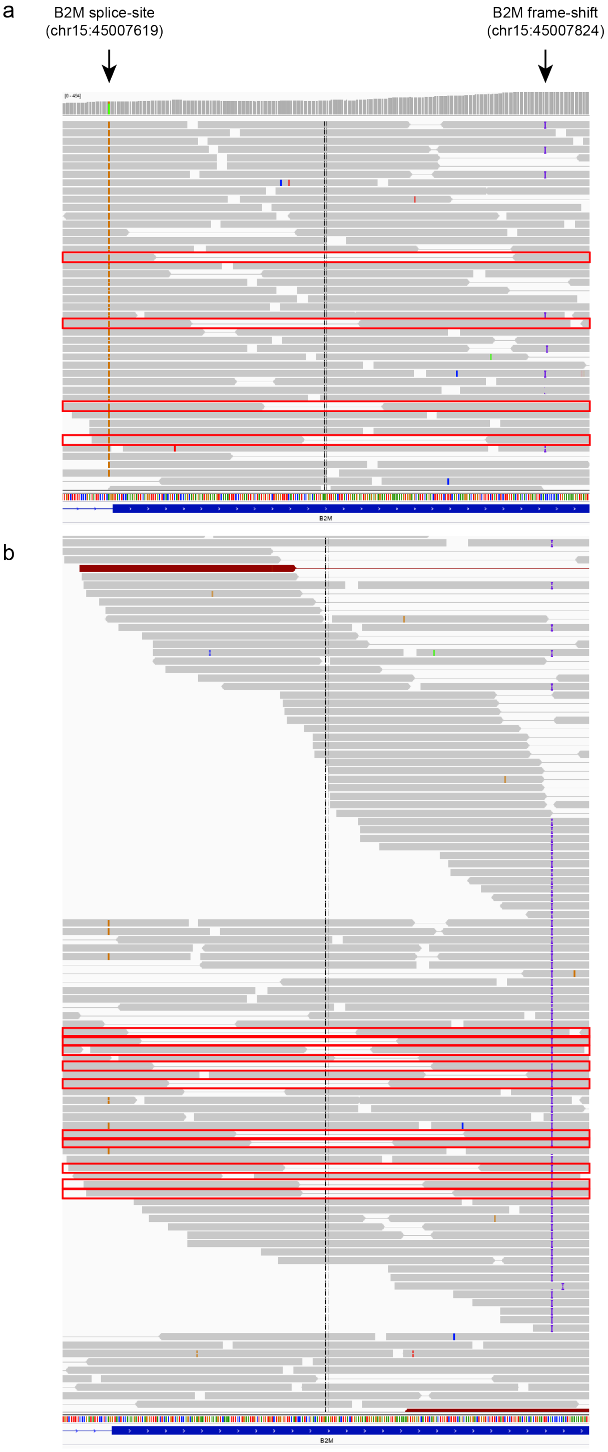

**Supplementary Figure 8. Trunk overestimation by single region analysis.**

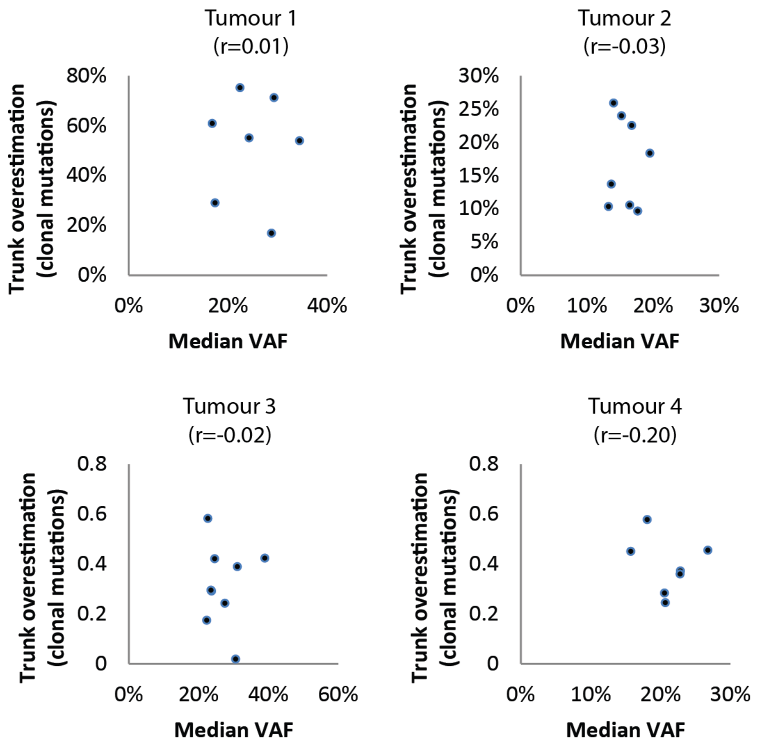

**Supplementary Figure 9. Mutation copy number of chromosome 8 in the STAD TCGA series.**

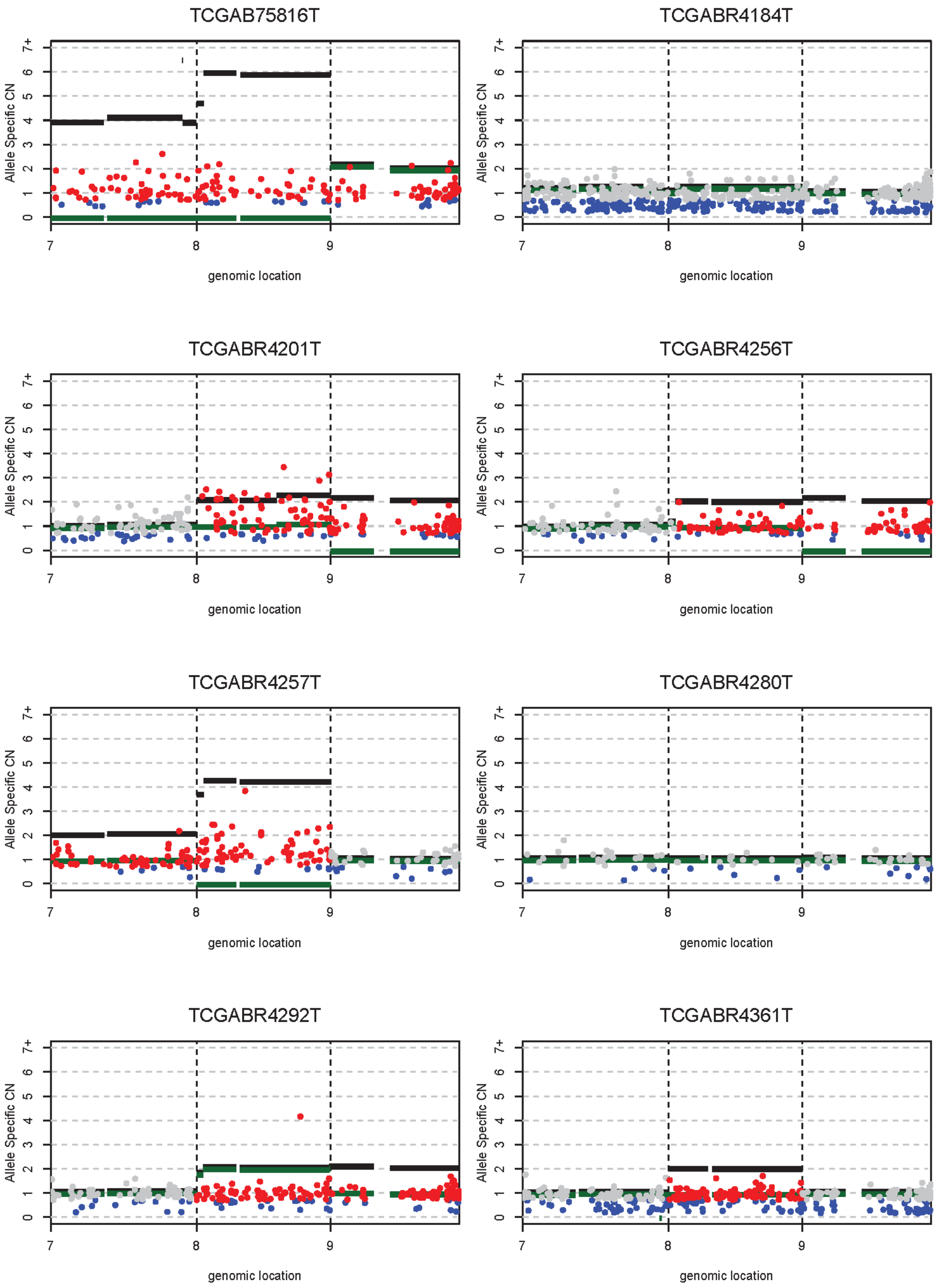

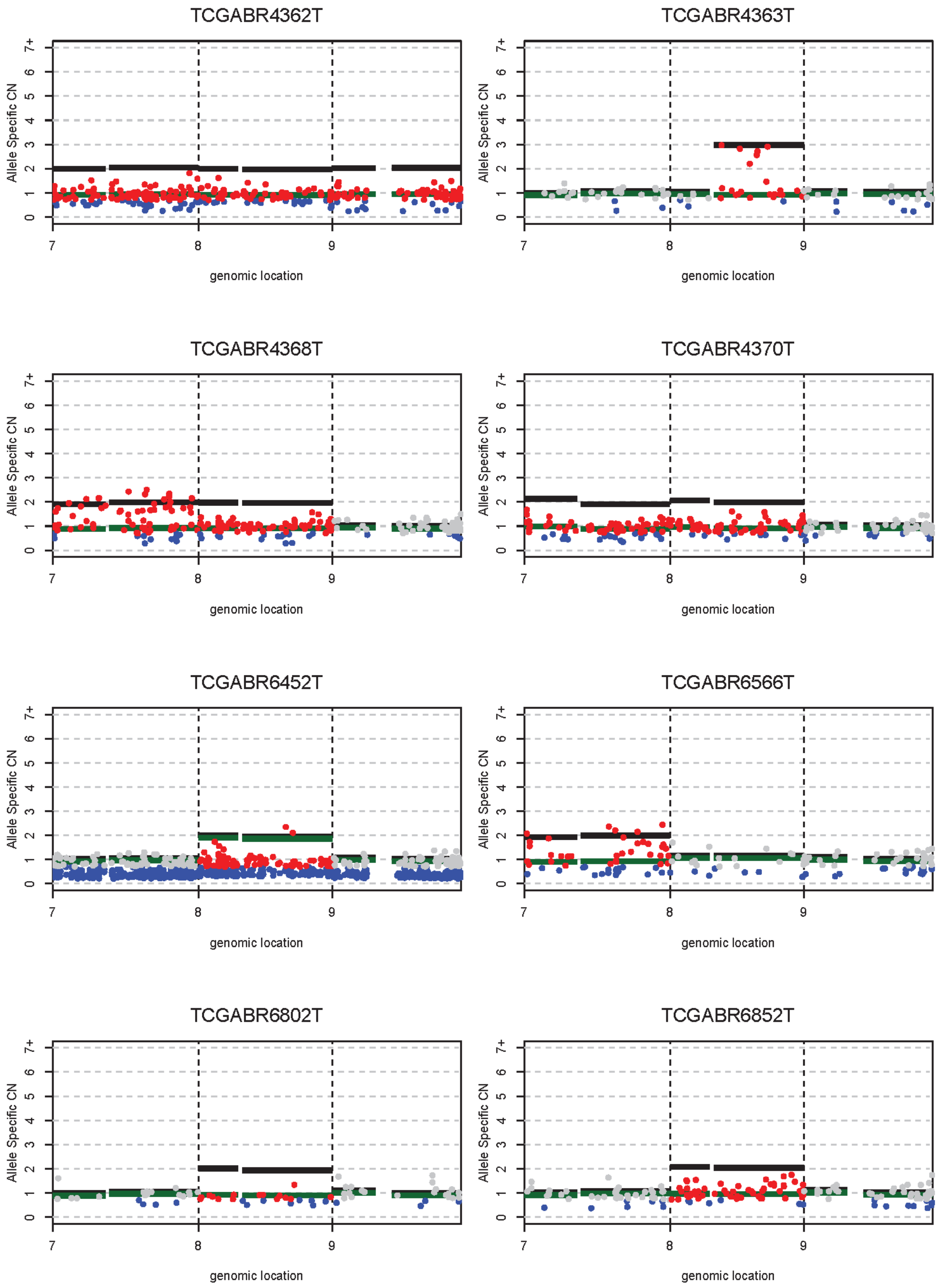
TCGABR4362T
TCGABR4363T
TCGABR4368T
TCGABR4370T
TCGABR6452T
TCGABR6566T
TCGABR6802T
TCGABR6852T
Allele Specific CN
0
1
2
3
4
5
6
7+
genomic location
7
8
9

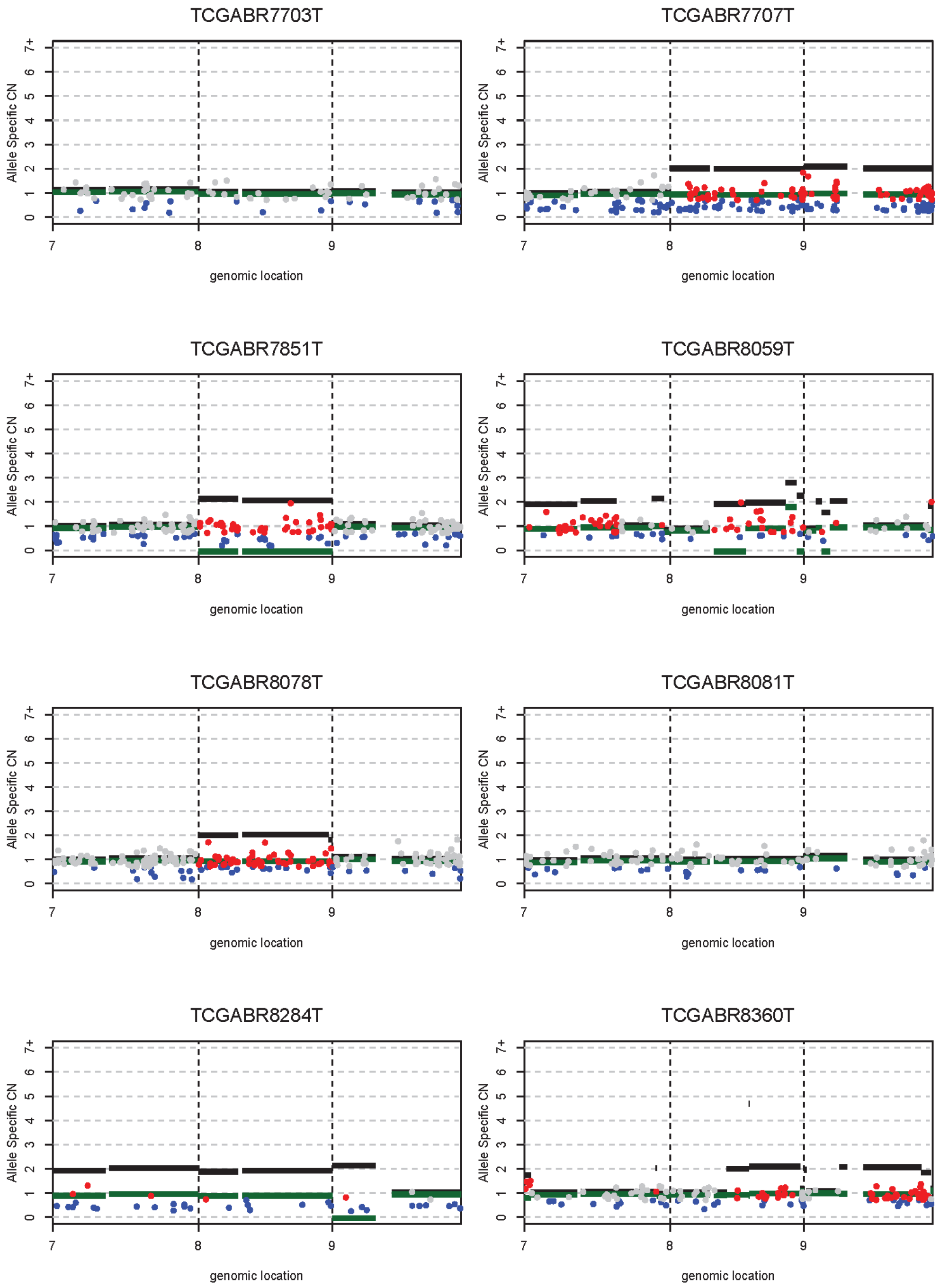
TCGABR7703T
TCGABR7707T
TCGABR7851T
TCGABR8059T
TCGABR8078T
TCGABR8081T
TCGABR8284T
TCGABR8360T
Allele Specific CN
0 1 2 3 4 5 6 7+
genomic location
7 8 9

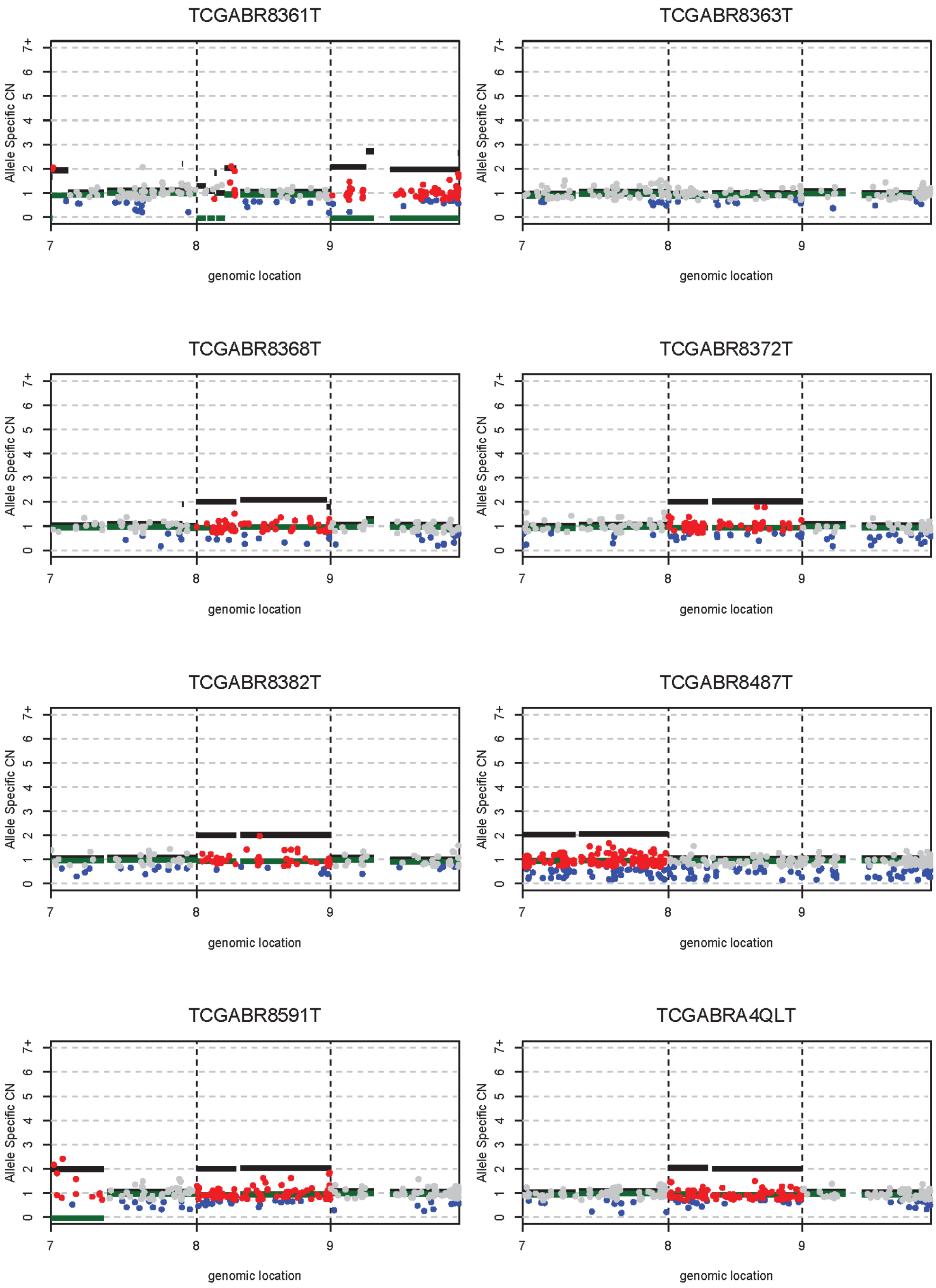
TCGABR8361T
TCGABR8363T
TCGABR8368T
TCGABR8372T
TCGABR8382T
TCGABR8487T
TCGABR8591T
TCGABRA4QLT
Allele Specific CN
0
1
2
3
4
5
6
7+
7
8
9
genomic location

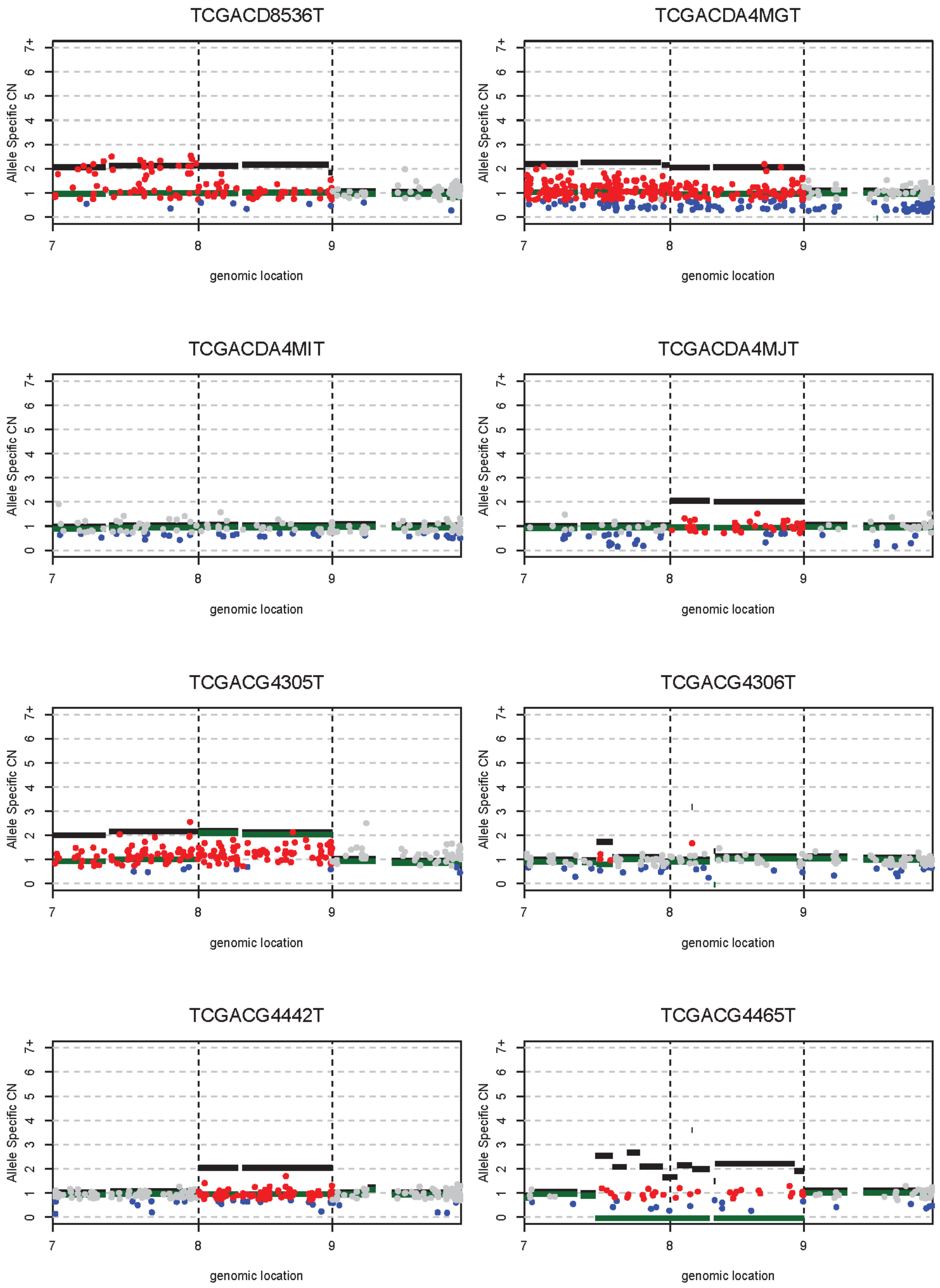
TCGACD8536T
TCGACDA4MGT
TCGACDA4MIT
TCGACDA4MJT
TCGACG4305T
TCGACG4306T
TCGACG4442T
TCGACG4465T
Allele Specific CN
0 1 2 3 4 5 6 7+
genomic location
7 8 9

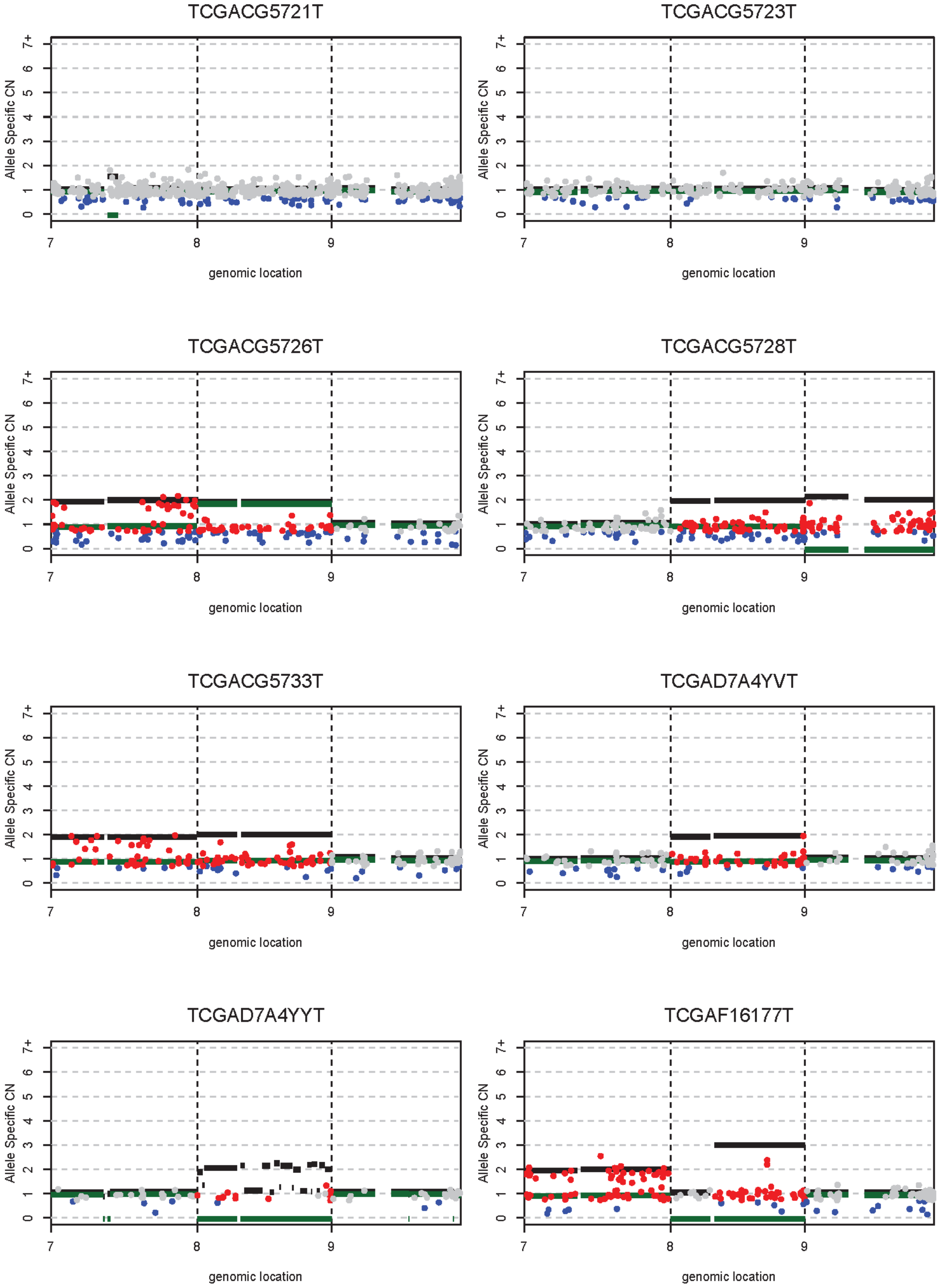
TCGACG5721T
TCGACG5723T
TCGACG5726T
TCGACG5728T
TCGACG5733T
TCGAD7A4YVT
TCGAD7A4YYT
TCGAF16177T
Allele Specific CN
0 1 2 3 4 5 6 7+
7 8 9
genomic location

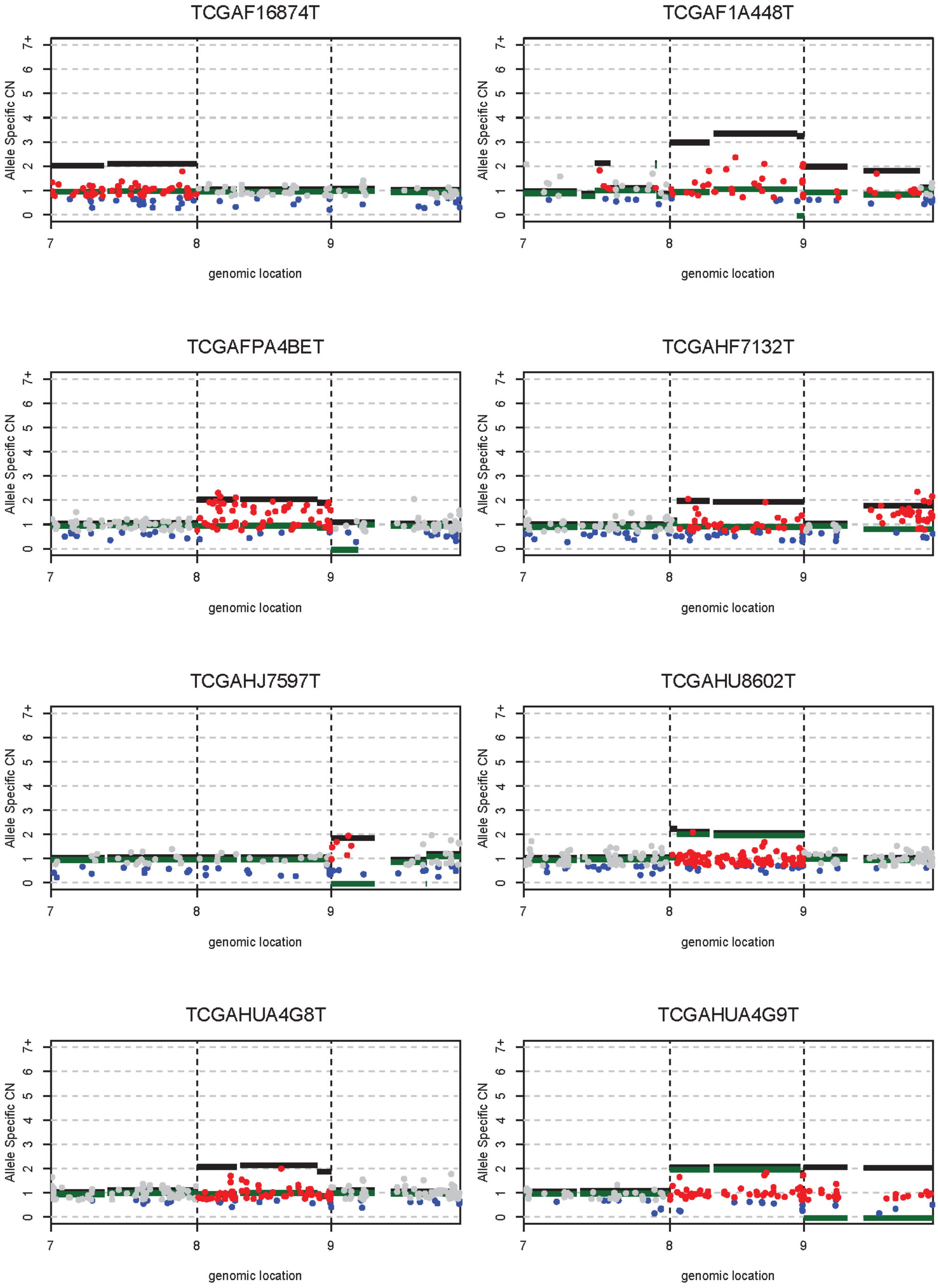
TCGAF16874T
TCGAF1A448T
TCGAFPA4BET
TCGAHF7132T
TCGAHJ7597T
TCGAHU8602T
TCGAHUA4G8T
TCGAHUA4G9T
Allele Specific CN
0 1 2 3 4 5 6 7+
genomic location
7 8 9

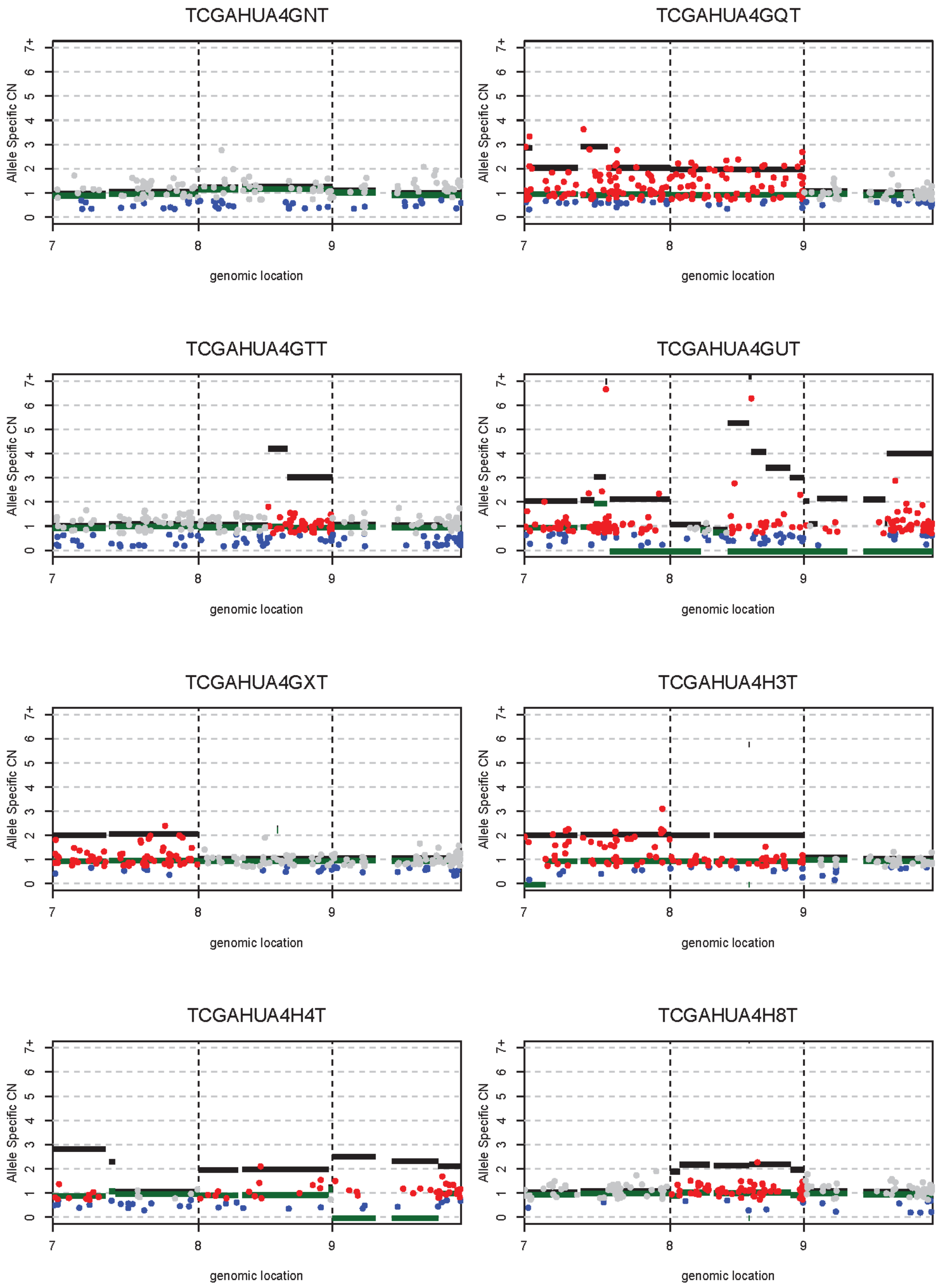
TCGAHUA4GNT
TCGAHUA4GQT
TCGAHUA4GTT
TCGAHUA4GUT
TCGAHUA4GXT
TCGAHUA4H3T
TCGAHUA4H4T
TCGAHUA4H8T
Allele Specific CN
genomic location
0
1
2
3
4
5
6
7+
7
8
9

**Supplementary Table 1. Mseq mutation calls and depths**

**Supplementary Table 2. Cell purity and ploidy**

**Supplementary Table 3. COSMIC signatures**

**Supplementary Table 4. Putative driver mutations**

**Supplementary Table 5. HLA mutation calls and LOH analysis**

**Supplementary Table 6. dN/dS**
